# Weak interactions drive selective proteome demixing and tune the differential response to environmental perturbations

**DOI:** 10.64898/2026.07.22.739830

**Authors:** Guido Narduzzi, Federico Zucca, Leo Willig, Ludovic Gillet, Ino Karemaker, Aleksander A Rebane, Alba Tomàs Sitjes, Leon Young, Moritz Becker, Simone Reber, Thomas Michaels, Paola Picotti, Gabriel E Neurohr

## Abstract

The intracellular space is a crowded environment where macromolecules perform distinct tasks despite pervasive “non-specific” interactions. Whether these interactions are functionally relevant and how they influence cellular organization remains unclear. Here, we developed QuPID-MS, which measures the propensity of proteins to phase separate in native cell extracts proteome-wide. We find that weak interactions drive condensation of half of the proteome in a crowding- and temperature-dependent manner and we present evidence that this proteome demixing occurs in cells. Importantly, protein condensation properties are conserved and broadly change when cells adapt to new environments, demonstrating that weak interactions are regulated and linked to function. Indeed, condensation of the growth regulator TORC1 coincides with its rapid inactivation, while high solubility of the stress-activated Hog1 ensures its activity across conditions. We thus uncover a fundamental organizing principle that allows tuning of cell growth to environmental fluctuations while ensuring other processes function robustly despite perturbations.

**Highlights:**

- Quantitative PEG induced demixing-MS (QuPID-MS) measures the tendency of proteins to condense in native cell extract.
- 53% of the yeast proteome is condensation-prone and demixes in a crowding- and temperature-dependent manner.
- The propensity to form condensates is globally modulated when cells adapt to new environmental conditions
- Weak interactions selectively organize the proteome in cells and differentially affect proteins involved in growth, signaling, and stress response.
- Dynamic proteome demixing allows coupling of macromolecule synthesis to environmental fluctuations while ensuring other processes function robustly despite perturbations.

**Graphical Abstract:** Proposed model for how selective proteome demixing tunes the differential cellular response to perturbations

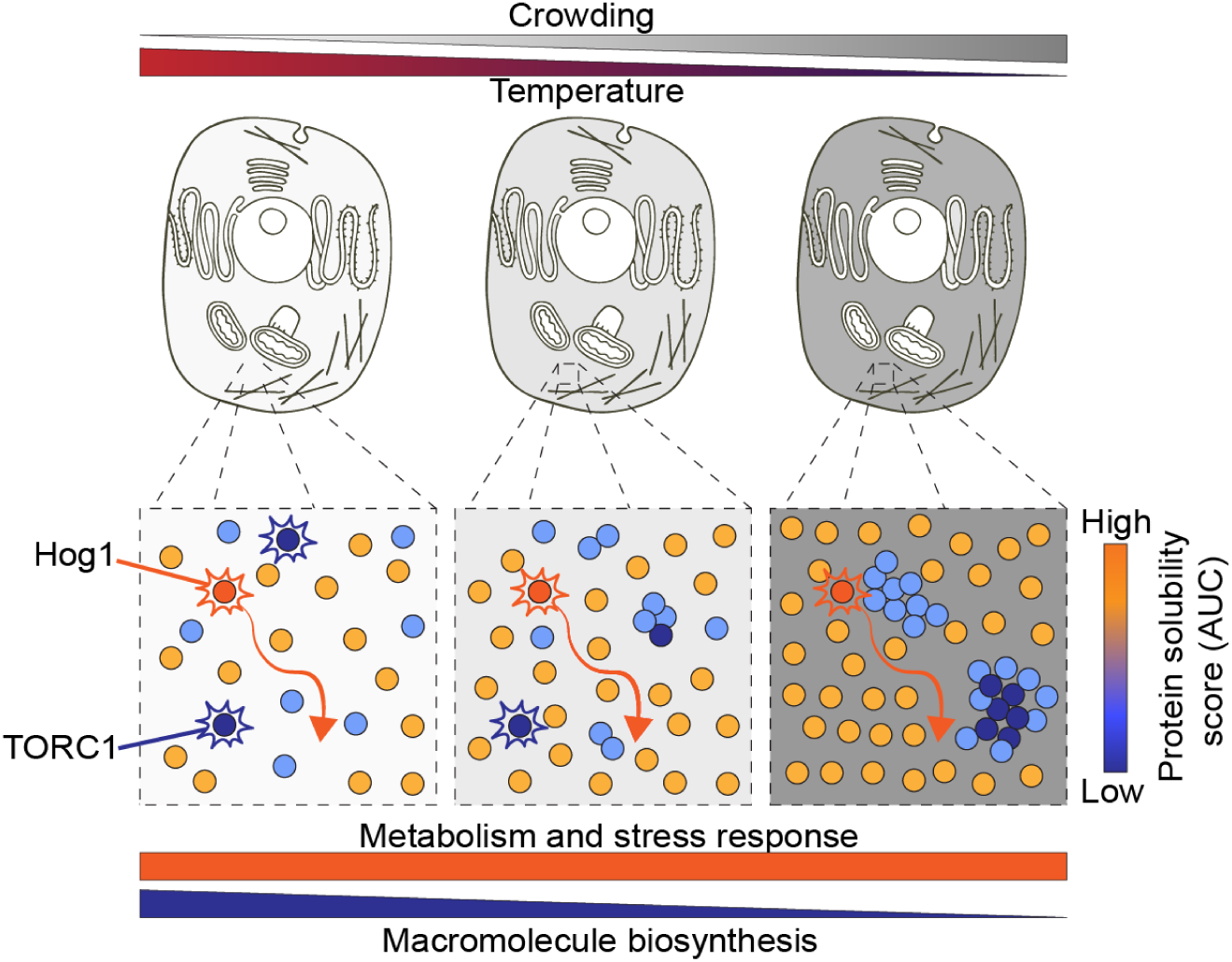

## INTRODUCTION

The inside of the cell is a highly crowded and complex environment where thousands of macromolecules perform countless distinct biochemical processes. How proteins can function in this packed environment is a mystery, especially because many proteins are sticky and interact promiscuously with many other macromolecules ^1–4^. Whether such “non-specific” weak interactions are functionally relevant and how they might influence the organization of the cell interior is unclear.

Weak interactions are particularly interesting because they are sensitive to changes in physico-chemical parameters such as temperature, ionic strength, and overall macromolecule concentration. Such changes occur in response to environmental fluctuations, during cell senescence, cell division and developmental processes, and they typically coincide with broad alterations in cell function ^5–13^.

At high protein concentrations, weak multivalent interactions can drive the formation of unstable protein assemblies ^14–19^. In a crowded environment, this process is further promoted by volume exclusion effects ^19,20^. Synthetic crowders such as polyethylene glycol (PEG) can therefore be used to drive the formation of such unstable conglomerates ^21^.

To understand how weak interactions might influence proteome organization, we treated native cell extract with PEG to induce the formation of protein assemblies that can be isolated by centrifugation. This allowed us to quantify the tendency of individual proteins to demix in a near native environment by mass spectrometry. Strikingly, we find that more than half of all proteins tend to form reversible assemblies in a crowding- and temperature-dependent manner, while other proteins remain fully soluble, and we provide evidence that this dynamic proteome organization occurs in cells. Importantly, we show that this protein feature is evolutionarily conserved, regulated during stress adaptation, and linked to cell function. Together, our data show that the cytoplasm is not a soup of randomly interacting sticky molecules but that proteins self-organize into dynamic assemblies that form or dissolve in response to physical changes. We propose that this dynamic demixing functionally divides the proteome into processes that respond sensitively to environmental changes and processes that continue to function robustly despite perturbations.

## RESULTS

### Quantitative PEG-induced demixing (QuPID) identifies condensation-prone proteins

In order to understand how weak interactions shape the organization of the proteome in cells, we generated minimally diluted, clear, native yeast cell extract ^13^ and added different concentrations of PEG (35kDa) to stabilize weak interaction-driven protein assemblies. Addition of PEG to extract induces the formation of microscopic droplets that are enriched for the P-body component Dcp2-GFP ^22^, which assembles into distinct foci (Figure 1A, S1A). Importantly, formation of these macroscopic and microscopic structures is fully reversible upon re-dilution of the PEG containing extract with undiluted extract or buffer (Figure S1A). While PEG partitions preferentially into the dilute fraction, proteins are enriched in the dense fraction (Figure S1B-E). Together these data demonstrate that this demixing process is not driven by co-condensation of PEG and proteins, but rather by weak, reversible interactions between the co-condensing molecules.

**Figure 1.**
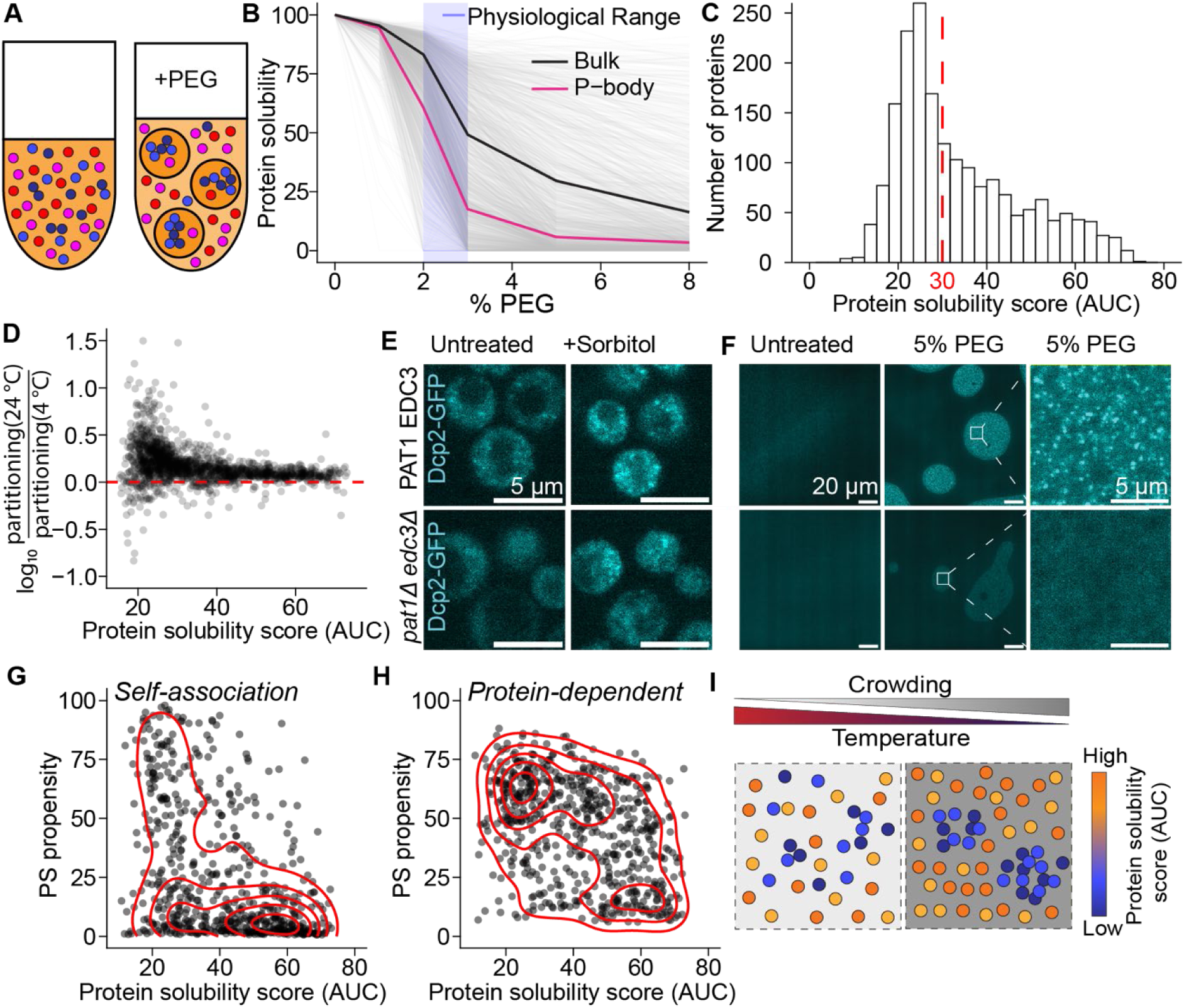
Weak interactions direct proteome self-organization. **(A)** Schematic of the quantitative PEG-induced demixing (QuPID) workflow: clear native cell extract is mixed with PEG35k, which leads to bulk phase-separation. The condensed and dilute fractions are separated by low-speed centrifugation and can be analyzed by mass spectrometry (QuPID-MS). **(B)** QuPID-MS was performed on yeast cell extract and the mass fraction of each protein remaining in the dilute fraction was determined at each concentration of PEG (thin grey lines). The bulk protein mass partitioning and the average of annotated P-body proteins are shown (bold lines). The putative range of crowding in cells is indicated in blue (see text). **(C)** The area under the curve (AUC) in B was calculated for each protein and the proteome wide distribution of this value plotted. The threshold at 30 is selected arbitrarily. **(D)** PEG induced fractionation was performed at 4 °C and 24 °C in yeast extract and the relative change in partitioning to the dilute fraction (mass in dilute over mass in dense fractions) is plotted against protein solubility scores at 24 °C for all proteins. **(E)** Images of yeast cells expressing the P-body component Dcp2-GFP before and 20 min after hyper-osmotic shock (1M Sorbitol) in wt and P-body deficient *pat1Δedc3Δ* mutant cells. **(F)** Images of yeast extract generated from the strains in (E), before and after PEG addition. **(G-H)** Comparison of our experimental measurements of protein solubility (C) with predictions from the PhaSePred algorithm ^25^ for phase separation propensity driven by self-association (G) or by association with other proteins (H). **(I)** Model of how weak interaction might drive environment dependent proteome de-mixing. Red lines indicate isodensity contours obtained from a bivariate kernel density estimate in G and H.

To determine the propensity of individual proteins to demix upon crowding changes, we separated the dilute and dense fractions by centrifugation and measured the partitioning of proteins by label-free data-independent acquisition (DIA) mass spectrometry. We named this workflow quantitative PEG-induced demixing-MS (QuPID-MS). This analysis revealed that proteins respond differently to crowding changes. While some proteins partition fully to the dense fraction at low PEG concentrations (3% PEG), other proteins remain fully soluble even at 8% PEG (Figure 1B). Components of stable complexes generally demix at the same PEG concentration (Figure S2A), showing that they remain intact during extract generation. We quantified the propensity of individual proteins to demix by integrating the area under the curve (AUC) in Figure 1B and will refer to this value as protein solubility score (Figure 1C, Table S1). This analysis was performed for 3233 proteins, which together account for almost 90% of protein mass in yeast ^23^, and it revealed that more than half (53%) of all proteins belong to a peak of proteins with a low solubility score (<30). Together, these poorly soluble proteins account for 43% of the mass of detected proteins.

If the formation of PEG induced protein assemblies is mainly driven by enthalpic attractive interactions between components of the extract, it should be enhanced at lower temperatures ^24^. Indeed, when we prepared QuPID-MS samples at 4 °C and 24 °C, almost all proteins showed an increased partitioning to the dense fraction at the lower temperature, but especially those that have a low solubility score (Figure 1D). This shows that demixing is driven by attractive interactions between macromolecules and protein partitioning is indicative of the strength of these interactions. Therefore, QuPID-MS can be used to estimate the interaction strength between proteins and their cellular matrix.

Analysis of protein features that underlie protein solubility in our assay revealed that membrane association, large size, positive charge, disordered regions and multiple folded domains all promote partitioning to the dense phase (Figure S2B-E). These are all features known to drive phase separation ^26,27^. Consistently, we find components of known biomolecular condensates, such as processing bodies (P-bodies) or the nucleolus, among the proteins that strongly demix at low PEG concentrations (Figure 1B, S2F-G). While nucleoli are constitutive condensates, P-bodies become brighter and more numerous upon osmotic compression ^28^ (Figure 1E). In our fractionation assay, P-body components transition from being soluble to partitioning mostly to the dense phase between 2% and 3% PEG (Figure 1B, S2F). We therefore estimate that the range of crowding proteins experience in cells corresponds to the region between 2% and 3% PEG in our assay.

When we generate yeast extract from cells that are unable to form P-bodies because they lack the scaffold proteins Pat1 and Edc3 ^28^ (Figure 1E), partitioning of the P-body client protein Dcp2-eGFP to the dense phase is reduced and it no longer forms punctate structures (Figure 1E-F). This shows that the same interactions that induce Dcp2 phase separation *in vivo* drive partitioning into the dense and dilute fractions after PEG-induced cytoplasm demixing.

Therefore, we compared our solubility score to *in silico* predictions of phase separation propensity. Indeed, there is a strong agreement between the predictions of the PhaSePred algorithm ^25^ and of protein solubility determined with QuPID-MS (Figure 1G-H). More specifically, we find that the prediction for protein-dependent phase separation correlates best with the propensity to demix (Figure 1H). This observation suggests that for most proteins, PEG induced demixing is driven by heterotypic interactions and follows the same molecular grammar that drives known examples of phase separation.

By quantifying how proteins demix upon PEG addition to cell extract we have therefore established a simple proteomic pipeline to measure the propensity of proteins to phase separate in the native cellular environment. Our measurements indicate that weak interactions drive large portions of the proteome into dynamic assemblies that form or dissolve in response to crowding and temperature changes while highly soluble proteins remain largely unaffected by such perturbations (Figure 1I).

### Condensation-prone proteins are immobilized in concentrated cell extract

While PEG is routinely used to induce phase separation *in vitro,* it is not a naturally occurring crowding regulator. Therefore, we wanted to test whether the solubility scores determined by QuPID (Figure 1C) inform about protein dynamics when modulating the overall extract concentration in the absence of PEG. To this end, we either diluted or concentrated extract using low molecular weight cutoff filters (<10 kDa) and the resulting flowthrough ^29^. This procedure allowed us to probe protein concentrations from 5mg/mL to 200mg/mL, exceeding the concentrations observed in unperturbed yeast cells (100 mg/mL ^30^). We then used fluorescence correlation spectroscopy (FCS) to measure the diffusion of representative GFP-tagged proteins at different overall protein concentrations. We selected cytoplasmic proteins that displayed either a low or high solubility score and that are not reported to be part of stable complexes. FCS measurements revealed that the diffusivity of condensation-prone proteins decreases rapidly with increasing total protein concentration, while the reduction was more gradual for highly soluble proteins (Figure 2A). Therefore, the propensity of proteins to condense upon PEG addition correlates with protein dynamics in concentrated native cell extract.

**Figure 2.**
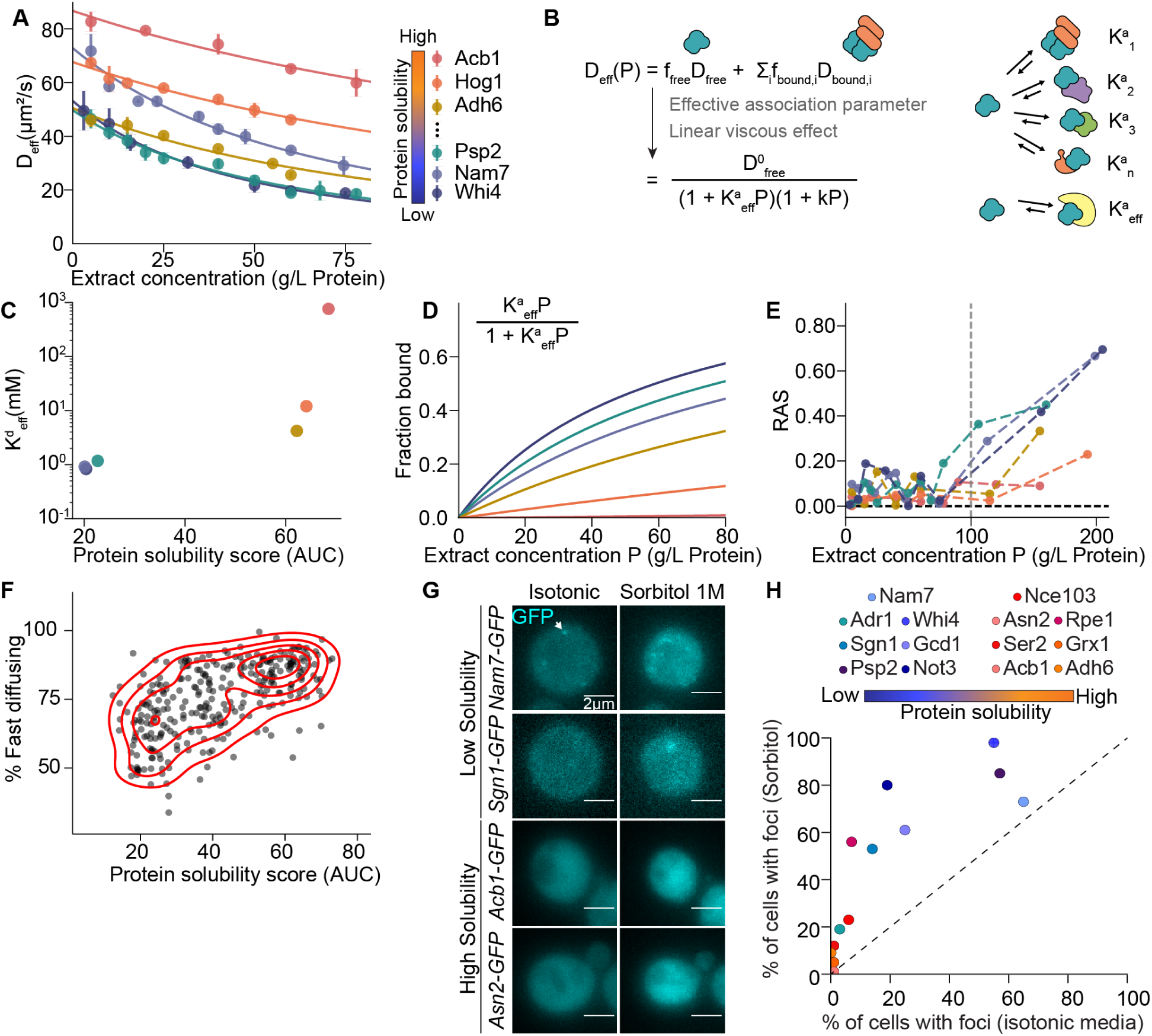
Weak interactions immobilize and spatially separate proteins in cells. **(A)** Extract of cells expressing GFP fusions of the indicated proteins were generated, concentrated using filtration and subsequently step-wise diluted to the indicated concentrations. Diffusion of the tagged proteins at different extract concentrations was measured by FCS and the effective diffusion coefficient D_eff_ was determined by fitting the data to a one-component FCS model. **(B)** Model used for fitting experimental data, assuming a global change in viscosity that affects all proteins equally and a protein-specific interaction strength with objects in the cytoplasm (see Supplementary Text). **(C)** The effective dissociation constant (K^d^_eff_) was extracted from the model in B and is plotted against the solubility score values from Figure 1C. **(D)** Model-derived estimates of the fraction of each protein that is bound to other cytoplasmic components as a function of overall extract concentration. **(E)** The quality of the one-component FCS fit to the raw data was evaluated using the Residual Asymmetry Score (RAS). At overall protein concentrations close to or higher than inside cells (dashed grey line) the one-component model is insufficient to describe the data for condensation-prone proteins but still works for soluble proteins. (**F**) Comparison of protein solubility scores and percentage of fast-diffusing protein obtained by ^31^ using in-cell FCS. Red lines indicate isodensity contours obtained from a bivariate kernel density estimate. **(G)** Confocal images of yeast cells expressing the indicated GFP-fusion proteins before and after 30 minutes treatment with 1M Sorbitol. **(H)** Quantification of (G) showing the percentage of cells expressing the indicated GFP fusion proteins containing at least 1 GFP focus. At least 100 cells were analyzed per condition.

Changes in viscosity alone should affect proteins similarly and therefore cannot explain the observed qualitative differences between condensation-prone and soluble proteins (Figure 2A, S3A). We therefore generated a quantitative model that also considers transient binding events to other cellular components (Figure 2B, Supplementary Text). Assuming that bound proteins diffuse significantly slower, we can use our model to determine the binding affinity between each protein and the intracellular environment (Figure 2C, S3B). Furthermore, it allows us to estimate the fraction of bound proteins at different extract concentrations. According to this model, highly soluble proteins remain mostly unbound even at high extract concentrations, whereas condensation-prone proteins undergo transient binding events that strongly impact their dynamics already at extract concentrations below those inside cells (Figure 2D). Consistent with this model, at concentrations close to or above the ones inside cells, a one-component autocorrelation-curve FCS model fails to describe the FCS data of condensation-prone proteins (Figure 2E). Together, these experiments show that condensation-prone proteins become increasingly bound and constrained when concentrating cell extract.

### Weak interactions organize the proteome in cells

Our *in vitro* experiments show that about half of all proteins tend to form crowding-sensitive assemblies in concentrated extracts. To determine whether this *in vitro* behavior is reflected in cells, we compared our proteomic data to orthogonal *in vivo* data sets. High throughput *in vivo* FCS measurements have been used to estimate the fractions of slow and fast diffusing protein pools in unperturbed yeast cells ^31^. The fraction of fast diffusing particles correlates with the solubility score of a protein (Figure 2F), showing that condensation-prone proteins are indeed partially sequestered in living cells. Furthermore, comparing this *in vivo* FCS dataset with our proteomic data confirmed that the crowding experienced by proteins in cells corresponds to yeast cell extract treated with 2%-3% PEG (Figure S3C).

To directly visualize how condensation-prone and soluble proteins respond to an increase in macromolecular concentration in yeast cells, we imaged cells expressing GFP-fused versions of selected proteins before and after hyperosmotic shock as previously performed ^32,33^. For these experiments, we chose representative proteins that appeared to have a diffuse cytoplasmic localization in unperturbed conditions ^34,35^. Visual inspection revealed that most of the imaged condensation-prone proteins already form small visible foci in unperturbed conditions and the fraction of cells containing such foci further increased after hyperosmotic shock (Figure 2G-H, S4). In contrast, selected soluble proteins rarely formed visible foci, even after hyperosmotic shocks. It is worth noting that these foci are much less prominent than, for example, P-bodies (Figure 1E), potentially reflecting their unstable and dynamic nature ^36^ and explaining why they have been overlooked by previous high throughput assays ^33^. Together, these observations show that large portions of the proteome are condensed and form distinct domains separated from the soluble proteome, even in unperturbed cells. When crowding increases this proteome demixing is exaggerated.

### Differential proteome demixing is a conserved organizing principle

Yeasts are exposed to frequent environmental changes such as temperature fluctuations and altered osmolarity. We wondered whether the described dynamic self-organization of the proteome is a unique feature of yeasts or a more general organizing principle also found in other organisms. We therefore determined whether the propensity to demix in a crowded environment is a conserved protein feature by performing QuPID-MS in extracts generated from HeLa cells (6684 quantified proteins) and *X. laevis* oocytes (5769 quantified proteins) at a single PEG concentration (Table S2). This experiment revealed that protein solubility is highly conserved among orthologs in these species (Figure 3A-B), indicating that weak interactions are under evolutionary pressure and critical for protein function.

**Figure 3.**
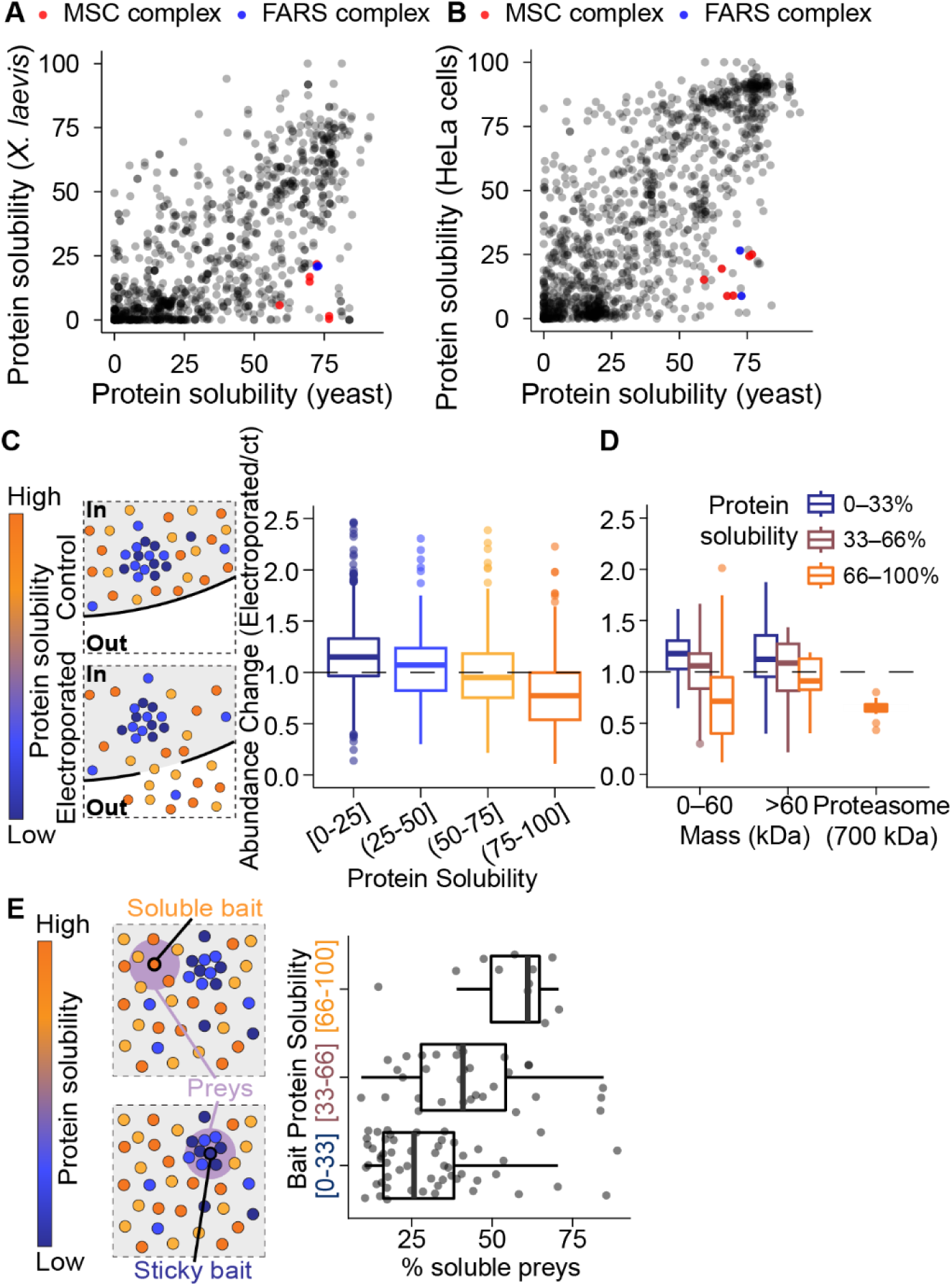
Weak interaction-driven proteome demixing is a conserved organizing principle. **(A, B)** Partitioning of orthologous proteins to the dilute phase in yeast, human (HeLa) and *X. laevis* egg cell extract treated with a PEG concentration that induces 40-60% bulk proteome condensation. The yeast-human comparison shows 1200 mapped ortholog pairs, the yeast-Xenopus comparison 995. **(C)** *Left:* Cartoon depicting preferential leakage of soluble proteins upon cell permeabilization. *Right:* Boxplot showing protein abundance in HEK293 cells before and after electroporation ^39^. Proteins are binned based on their propensity to condense in HeLa extract treated with PEG (Figure 3A). **(D)** Same data as in C but only shown for monomeric proteins split into the indicated size bins and for all detected proteasome subunits. **(E)** *Left:* Cartoon depicting in-cell proximity ligation proteomics and how it reports on the local environment of proteins. *Right:* Bait proteins of a large scale BioID study in human HEK293 cells ^40^ were binned according to their protein solubility. For each bait protein, the composition of interactors was analyzed and the percentage of soluble prey proteins (>33% in the dilute fraction) is plotted.

Importantly, there are exceptions to this rule. One example of this is a class of tRNA synthetases, which are soluble in yeast but condensation-prone in human and frog extracts (Figure 3A-B). Many of these enzymes are known to be part of the megadalton-sized multi synthetase complex, a supramolecular assembly that forms in human and frog cells but not in yeast ^37^. Interestingly, components of the phenylalanyl synthetase complex (FARS) also show the same behavior (Figure 3A-B). This complex is thought to exist as a much smaller tetrameric complex ^38^, but our data suggest that it might also be part of a large supramolecular assembly in the crowded environment of the cytoplasm. Overall, these experiments show that protein solubility is a highly conserved feature but our method can also detect differences in proteome organization between these species.

In yeast, condensation-prone proteins are partially immobilized in cells and spatially demix from soluble proteins. To test whether parts of the proteome are also immobilized in human cells, we investigated which proteins leak out of cells upon permeabilization of the membrane. For this purpose, we re-analyzed a previously generated dataset that carefully examined the intracellular proteome before and after electroporation in human HEK cells ^39^. Strikingly, only proteins determined to be highly soluble leak out of cells through electroporation-induced pores (Figure 3C-D). This effect is not driven by the smaller average size of soluble proteins. Amongst equally sized proteins only the most soluble proteins diffuse out of cells and even large soluble complexes, such as the proteasome, rapidly leak out (Figure 3D). Together, these data demonstrate that condensation-prone proteins are partially immobilized in human cells.

If condensation-prone proteins cluster together in cells this could lead to the formation of microcompartments enriched for either soluble or condensation-prone proteins. To test this idea, we compared our data with a large proximity labeling BioID dataset obtained in unperturbed human HEK293 cells ^40^. Indeed, condensation-prone bait proteins tend to interact more frequently with other condensation-prone proteins, while soluble bait proteins mostly interact with soluble ones (Figure 3E). Therefore, condensation-prone proteins are to some degree both immobilized and spatially removed from soluble proteins also in human cells. By evaluating protein solubility in multiple organisms, we therefore show that weak interactions underlie a conserved self-organization principle of the proteome.

### QuPID-MS can be used to study the regulation of condensate assembly

While weak-interaction-driven assemblies can respond directly to biophysical changes, in cells there are additional layers of regulation that might stabilize or destabilize such assemblies. We wondered whether QuPID-MS can be used to study changes in proteome self-organization after genetic perturbations.

The composition and assembly pathways of P-bodies in cells have been extensively studied ^22,41,42^. Deletion of the P-body scaffold proteins Pat1 and Edc3 for example impairs the recruitment of client proteins to P-bodies in cells (Figure 1E ^43^). To determine whether we can detect these changes using our proteomic approach, we induced proteome demixing in extracts from *wild type* and *pat1Δ edc3Δ* double mutant cells using 3% PEG and compared proteome partitioning. Indeed, many P-body proteins ^42^ become less condensation-prone when the scaffold proteins are absent (Figure 4A, Table S1). The increase in solubility correlates with the reported *in vivo* enrichment of proteins into P-bodies ^42^ (Figure 4B), demonstrating that our assay can quantitatively capture P-body composition. Strikingly, this relationship is not observed for components that are shared between P-bodies and stress granules, a related cytoplasmic condensate with a set of overlapping components ^44^ (Figure 4B). Together, these data show that QuPID-MS can be used to study the regulation of condensate assembly and they also highlight that this is a complex process because condensates are heterogenous and can share common components.

**Figure 4.**
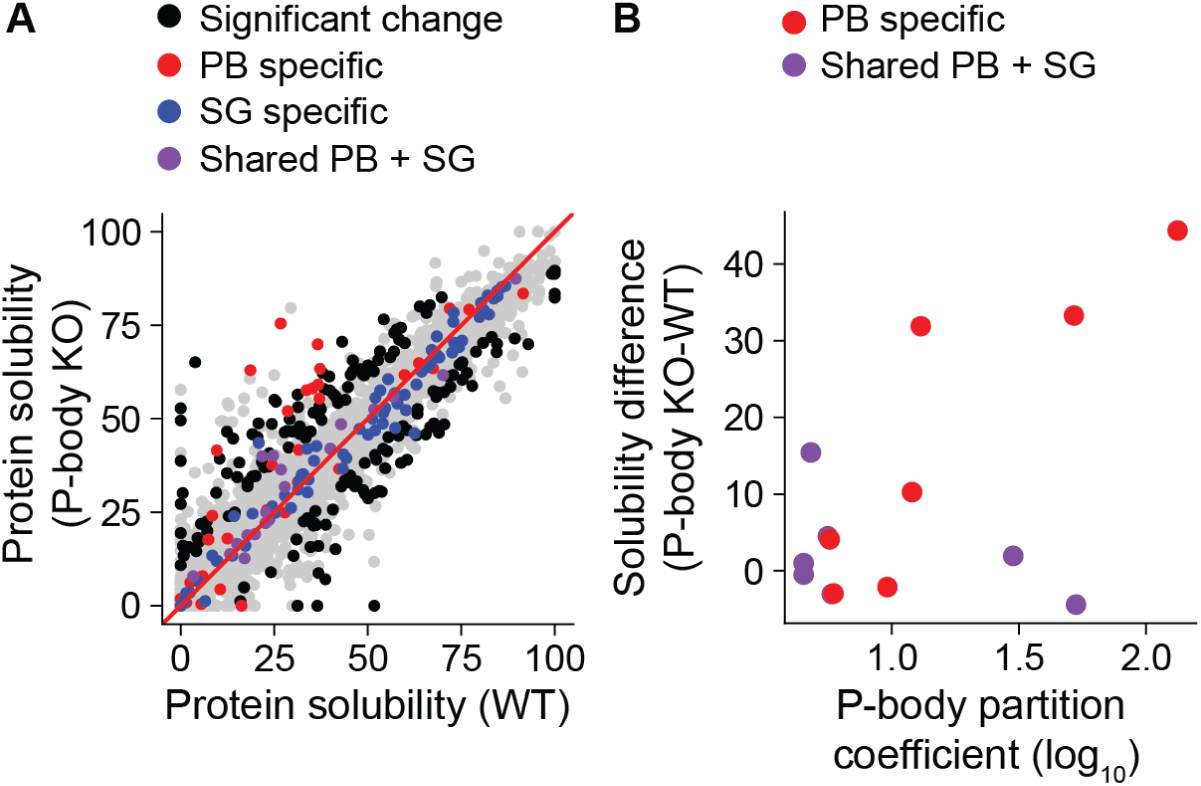
QuPID-MS reveals changes in P-body composition upon removal of scaffold proteins. **(A)** Comparison of protein solubility at a fixed PEG concentration in yeast extract from a WT and a *pat1Δ edc3Δ* mutant strain *(P-body KO)*, which is defective in P-body formation *in vivo*. Protein solubility corresponds to the percentage of total protein partitioning to the dilute phase after addition of 3% PEG. Components of P-bodies (PB) and stress granules (SG) are colored, other proteins with significantly changed solubility between the conditions are shown in black. **(B)** Reported *in vivo* partition coefficients of individual proteins to the P-body^42^ (x-axis) plotted against the difference in solubility in extracts of WT and P-body KO strains from the data in (A). Both P-body specific components and components shared between P-bodies and stress granules are shown.

### Proteome demixing is regulated when cells adapt to stress

Environmental perturbations such as changes in nutrient availability induce proteome rearrangements that lead to the formation of large supramolecular assemblies and filaments ^45–50^. Assembly of these structures has been proposed to help cells adapt to the new conditions ^48,51^. We wondered whether QuPID-MS can be used to characterize this altered proteome configuration in starved cells. To address this point, we generated extracts from exponentially growing cultures and from cultures that ran out of nutrients and reached stationary phase. Of note, while intracellular pH drops in starved cells ^52^ we adjusted extract pH to 7.1 prior to extract demixing for consistency with other experiments. QuPID-MS indeed revealed striking global shifts in protein solubility that affected 28% of all detected proteins (Figure 5A).

**Figure 5.**
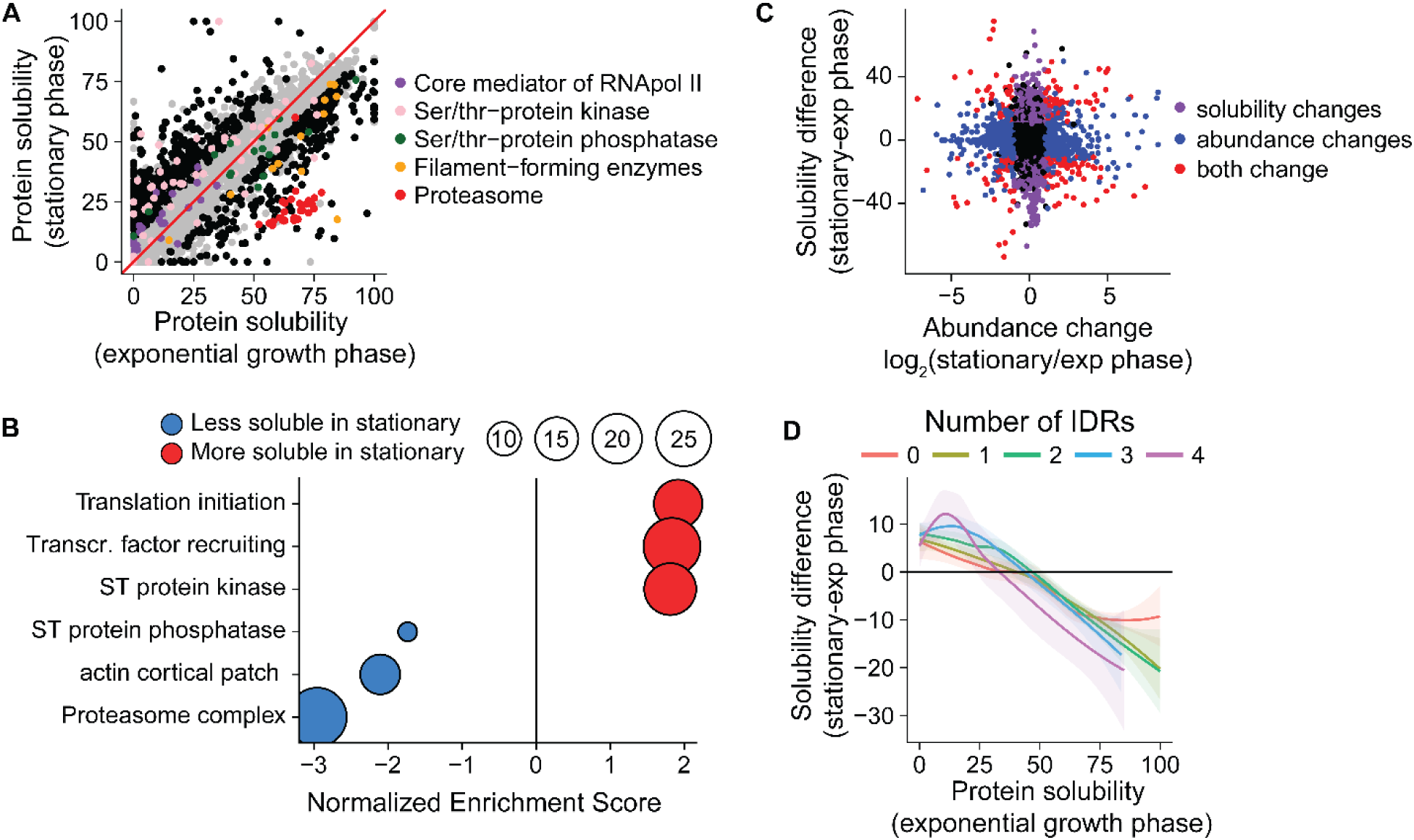
Starvation alters condensation propensity in a global and protein-specific manner. **(A)** Comparison of protein solubility at a fixed PEG concentration in yeast extracts from a WT strain collected in either exponentially growing cultures or from cultures that were grown for 24h and reached early stationary phase. pH was adjusted to 7.1 in both extracts prior to PEG addition. Components of selected complexes and protein classes are colored. Proteins with significantly changing solubility between the conditions that are not part of these groups are shown in black. **(B)** Proteins were ranked based on the difference in solubility as cells transition from exponential to stationary phase and GSEA analysis was performed. Select groups of proteins that either become more or less soluble during this transition are shown. **(C)** Comparison of changes in protein abundance and altered protein solubility of the experiment shown in (A). **(D)** Loess regression of the difference in protein solubility between the two conditions from A plotted against initial protein solubility. The regression was performed independently for protein groups with a given number of intrinsically disordered regions (IDRs). Shaded areas show the relative pointwise 95% confidence interval of the regression.

Firstly, many highly soluble proteins become less soluble in extract from stationary phase cultures. Gene set enrichment analysis revealed that amongst these proteins, components of the proteasome and cortical actin patches are highly enriched (Figure 5A-B). This reduction in solubility likely reflects previously described observations. The proteasome for example assembles into storage granules as cells enter starvation ^49,50^ and dynamic cortical actin patches are known to become more stable ^53^. Similarly, many metabolic enzymes known to form filamentous structures during starvation ^47,48,54^ become less soluble in extracts from starved cells (Figure 5A). QuPID-MS is therefore able to detect known proteome rearrangements when cells adapt to changing environments. Importantly, our data suggest that these previously observed changes are part of a global proteome-wide re-organization that renders many soluble proteins more condensation-prone during the transition to stationary phase.

Secondly, and more surprisingly, many condensation-prone proteins tend to become more soluble when cells enter stationary phase (Figure 5A). Components of the core transcription- and translation initiation machinery for example become more solubilized in starved cell extract, potentially reflecting the reduced biosynthetic activity (Figure 5B).

Interestingly, while protein kinases tend to become more soluble in stationary phase, protein phosphatases become more condensation prone. This differential solubility change might contribute to a global shift in the balance between kinases and phosphatases as cells enter stationary phase.

We next wondered what underlies these changes in the condensation propensity of proteins. Proteome composition changes dramatically as cells enter stationary phase and it has been proposed that many proteins are expressed near their solubility limit ^55,56^. We therefore hypothesized that changes in protein abundance could be responsible for the altered protein solubility. To test this idea, we quantified the protein abundance in extracts from growing and starved cells. These protein abundance changes, however, do not correlate with the changes in protein solubility (Figure 5C), demonstrating that protein solubility is not simply determined by protein expression level but that there are additional levels of regulation.

We therefore analyzed whether specific protein features might drive the change in protein solubility upon entry into starvation. Our initial analysis revealed that the presence of intrinsically disordered protein regions biases proteins to be condensation-prone. As condensation-prone proteins globally tend to become more soluble when cells enter stationary phase, IDR containing proteins are also overall more likely to become solubilized. But even if we take the initial protein solubility into account, IDR containing proteins still tend to become more solubilized than proteins not containing IDRs (Figure 5D). The presence of IDRs in highly soluble proteins on the other hand increased their tendency to become condensation-prone. The presence of disordered regions therefore predisposes proteins to change solubility when cells enter stationary phase. Importantly, protein disorder cannot explain all solubility changes as cells enter stationary phase, suggesting that there are additional regulatory mechanisms. Overall, these observations show that weak interaction driven proteome organization is subject to regulation when cells adapt to changing environments, suggesting that they are relevant for cell physiology.

### Differential proteome demixing tunes the responses to environmental perturbations

To identify potential functional implications of organizing the proteome into soluble and condensed phases, we performed a Gene Set Enrichment Analysis (GSEA) on protein solubility scores. This revealed that proteins involved in related biological processes have similar condensation properties (Table S2). Metabolic enzymes and proteins that are part of cellular stress response pathways tend to remain soluble even at high crowding levels (Figure 6A), whereas proteins involved in regulating transcription and protein synthesis tend to become immobilized upon increased crowding. As the propensity to condense is a conserved protein feature, this correlation between protein solubility and protein function can also be observed for human and *X. laevis* proteins (Figure 6B, S5A-B, Table S2). The dynamic demixing of the proteome in response to crowding changes therefore might allow tuning of certain cellular processes while leaving others unchanged.

**Figure 6.**
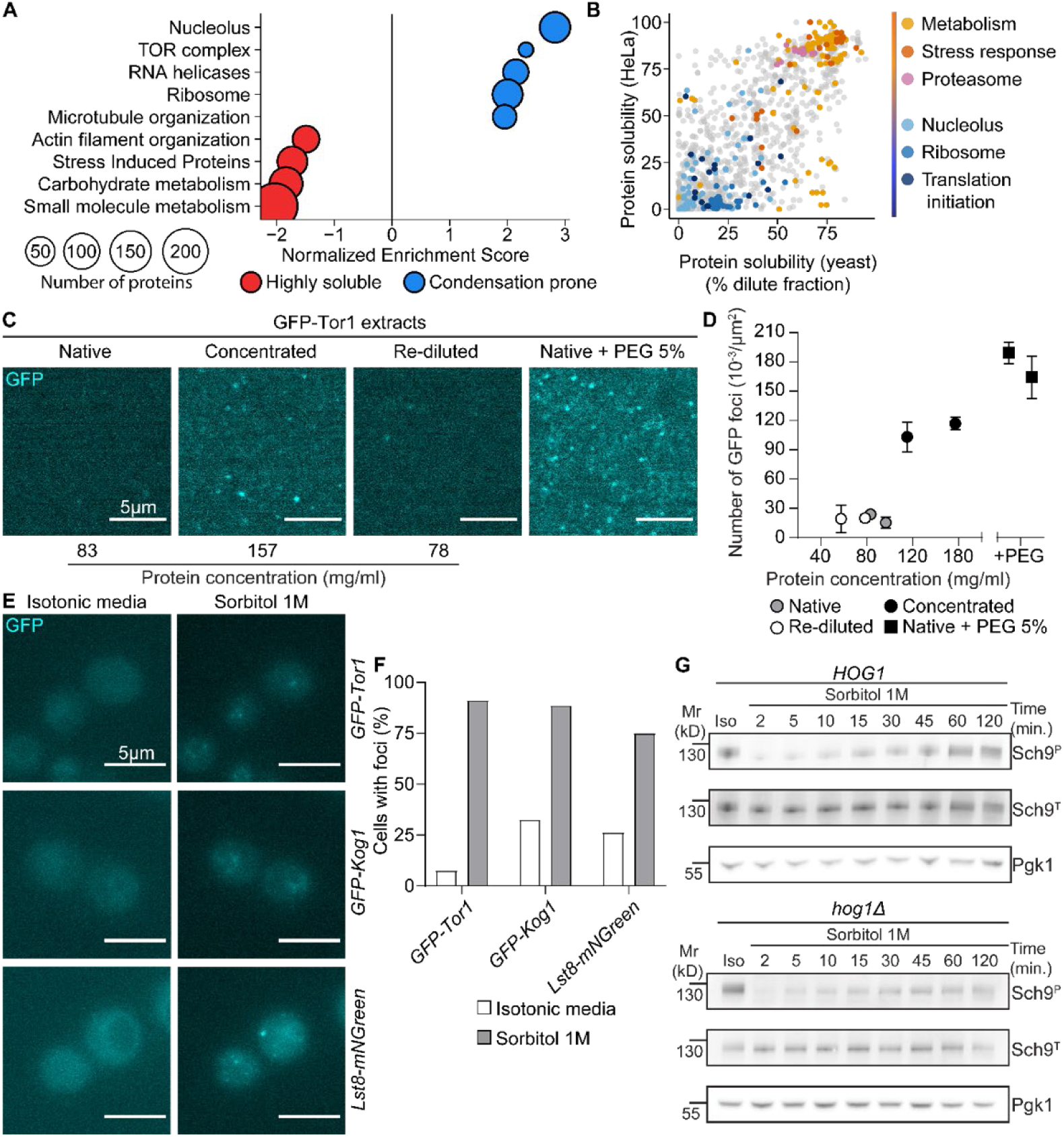
Protein condensation couples protein function to environmental fluctuations. **(A)** Yeast proteins were ranked based on their solubility score and a single sample GSEA projection was performed. Selected sets of proteins enriched for either soluble (red) or condensation-prone proteins are shown. **(B)** Solubility for orthologous yeast and human proteins as shown in Figure 3A but with conserved functional modules highlighted. **(C)** Images of yeast extracts of cells expressing GFP-Tor1 right after the extraction (Native), upon concentration (Concentrated), re-diluted after the concentration (Re-diluted), or mixed with PEG at the final concentration of 5% (Native + PEG 5%; the image shown is of the dense fraction). **(D)** Quantification of the number of GFP-Tor1 foci in the samples treated as in (C). **(E)** Images of yeast cells expressing the TORC1 subunits *GFP-Tor1, GFP-Kog1* and *Lst8-mNGreen* before and 30 minutes after Sorbitol addition. **(F)** Quantification of the number of cells containing GFP-foci from cells treated as in (E). At least 100 cells were analyzed per condition. **(G)** Time course analysis of the indicated yeast cells treated with Sorbitol for the indicated times. Western blot for phosphorylated TORC1 substrate Sch9-S758-P (TORC1 target), total Sch9 and PGK1 (loading control). The experiment was repeated 2 times with similar results.

Protein synthesis for example has been shown to be coupled to macromolecular crowding in bacteria, budding yeast, human cells and *Xenopus* extracts ^29,57–59^. In human cells, this coupling depends on the activity of the central growth regulator mTOR complex 1 (mTORC1) ^57^. Intriguingly, we find TOR complex 1 to form reversible condensates upon increased crowding in cell extract treated with PEG or when we concentrate extracts by filtration (Figure 6C-D). Visible TORC1 foci can also be observed in yeast cells after hyperosmotic shocks (Figure 6E-F), which also cause a rapid inactivation of TORC1 as previously reported ^60^ (Figure 6G). Importantly, this rapid TORC1 inactivation is independent of Hog1, the osmo-stress induced MAP kinase and p38 homolog in yeast (Figure 6G). Together, these data show that crowding-induced TORC1 condensation correlates with rapid TORC1 inactivation that occurs independent of stress signaling.

In contrast to translation regulators, proteins needed for cellular stress responses tend to remain soluble in over-crowded conditions (Figure 6A). Retention of solubility could make these processes robust to environmental perturbations. Hog1 for example remains soluble even at high crowding (Figure 7A-B). Upon hyperosmotic shock, Hog1 translocates to the nucleus to mount a transcriptional response, and its high solubility might be a key feature to maintain it mobile in highly crowed conditions ^61^. To test this idea, we fused Hog1 to the low complexity domain of the condensation-prone DEAD-box helicase Dpb1 ^62^. This chimera protein partitions more to the dense fraction in PEG-induced condensates (Figure 7A-B) and its diffusion is rapidly slowed down upon extract concentration (Figure 7C), showing that we successfully generated a condensation-prone allele of Hog1. While hyperosmotic shock still leads to a rapid phosphorylation of Hog1-Dpd1-LCD (Figure 7D), translocation of this protein to the nucleus is severely impaired (Figure 7E-F). Consistent with this observation, the Hog1 dependent transcriptional response is impaired in cells expressing the endogenously tagged Hog1-Dpd1-LCD (Figure 7G). This demonstrates that the absence of weak interactions between Hog1 and the cytoplasmic matrix is crucial for its function in an overcrowded environment.

**Figure 7.**
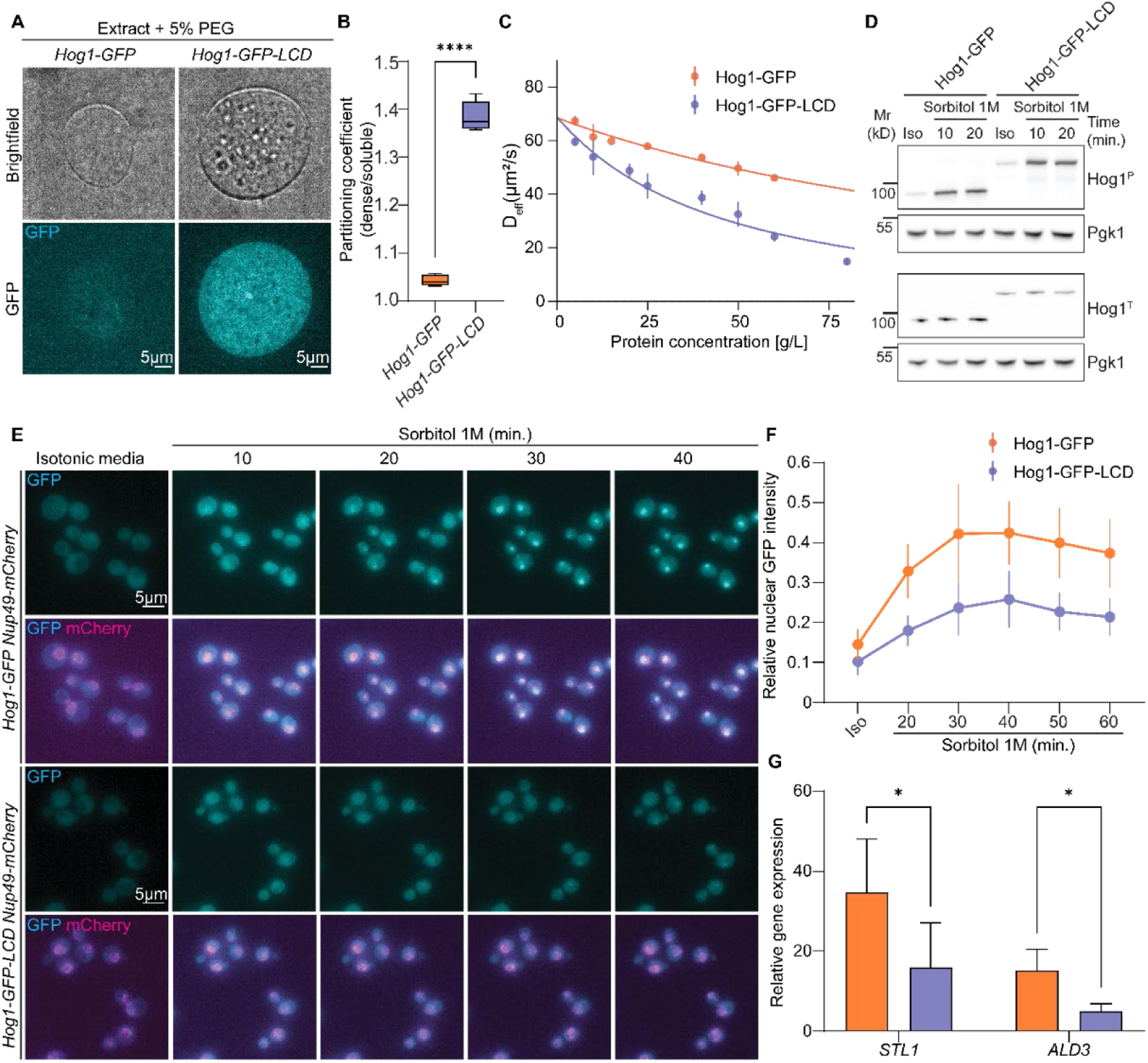
High Hog1 solubility allows efficient stress signaling in crowded conditions. **(A)** Yeast extracts of cells expressing Hog1-GFP or Hog1-GFP-LCD imaged after addition of 5% PEG. **(B)** Quantification of the partitioning coefficient of the chimera proteins from A to the dilute and dense fraction. The experiment was repeated 3 times with similar results. **(C)** Extract of cells expressing Hog1-GFP or Hog1-GFP-LCD were generated and diluted to the indicated concentrations. Diffusion coefficient D_eff_ of the tagged proteins was determined using FCS and by using one-component fitting of the data. **(D)** Western blot analysis of Hog1 phosphorylation during Sorbitol treatment using a phospho-p38 (Thr180/Tyr182) antibody. **(E)** Live cell imaging of cells with the indicated genotypes before and after Sorbitol addition. **(F)** Quantification of relative nuclear GFP intensity from (E). At least 100 cells were analyzed per condition. **(G)** Expression of the Hog1 target genes STL1 and ALD3 was determined by RT-qPCR in cells expressing Hog1-GFP (orange) or Hog1-GFP-LCD (blue) before and 30 minutes after Sorbitol addition. The graph shows the gene expression relative to ACT1 and normalized to the expression in isotonic conditions. N=3.

## DISCUSSION

We developed a proteome-wide assay to measure the propensity of proteins to condense in their native environment. Our data show that this tendency is driven by weak interactions and that it underlies the self-organization of more than half of the proteome into unstable assemblies that form or dissolve in response to changes in crowding and temperature. This propensity is strongly conserved, regulated during stress adaptation and it is linked to biological function. We show that crowding dependent condensation of TORC1, a master regulator of growth and biosynthesis, correlates with its rapid inactivation. The high solubility of the Hog1 MAP kinase, on the other hand, is needed for efficient stress signaling in overcrowded conditions (Figure 7).

### A new method to study unstable supramolecular assemblies

We used PEG induced extract fractionation to measure the tendency of proteins to reversibly assemble with other cytoplasmic components. Thanks to the use of PEG, we can measure this tendency at physiological crowding levels and even in slightly dilute extracts. In addition, the approach is technically simple, cheap, and therefore widely applicable, allowing comparable systematic studies on the composition and regulation of unstable condensates in different biological systems. As a proof of principle, we use this approach to measure how strongly the P-body scaffold proteins Pat1 and Edc3 contribute to recruiting different P-body components.

Previous systematic approaches to identify condensates that use diluted extract and detergents identified only 5-10% of all proteins to be condensation-prone ^63–65^, which was at odds with in silico predictions that estimated 40-80% of proteins to phase separate ^66^. This highlights the importance of measuring labile condensates in native conditions and across the physiological crowding range. A recent study, using undiluted Xenopus extracts and a filtration-based approach, identified 18% of proteins to be condensation-prone ^67^, still significantly less than what we observe with our method. This discrepancy may result from the fact that the filtration-based approach is unable to detect micro-condensates smaller than the used pore size or that it focuses on detecting assemblies with liquid properties. Despite using a completely different method, at the individual protein level our measurements in frog extract correlate well with their results (Figure S6A). More surprisingly, we also found a strong correlation between our protein solubility score and the ability for proteins to re-solubilize after complete dehydration of yeast extract ^68^ (Figure S6B). This suggests that a similar underlying biophysical mechanism drives reversible condensation and irreversible aggregation in the proteome depending on the severity of the perturbation.

### Weak interactions organize the cell interior

Biomolecular condensates have been recognized to compartmentalize the cell interior since their discovery ^17^, allowing cells to dynamically respond to environmental changes ^18,41,69,70^. Our measurements show that these specific condensates might only be the tip of the iceberg, reflecting a more general, underlying organizing principle that involves the majority of proteins.

This bulk protein condensation might spatially compartmentalize the cell interior, directly affecting the mobility and transport of large structures. The idea that parts of the proteome form an amorphous network has been proposed by Keith Porter in the 1970s based on electron micrographs, but the idea was later discarded because of technical concerns ^71^. More recent evidence in live cells however supports the idea that the cytoplasm is compartmentalized, impacting the diffusion of small proteins and large particles, and that this ultrastructure dynamically changes in response to osmotic perturbations ^7–9,72,73^.

Condensation of large fractions of the proteome has also been suggested to influence intracellular properties such as the water potential and ion homeostasis ^74,75^. We propose that in addition, bulk proteome condensation could free up space for highly soluble proteins and thereby buffer their concentration and ensure continued mobility. This effect could be amplified by the fact that protein concentration in the condensed cytoplasmic phase can increase without a corresponding volume increase (Figure S1C-E). Such a condensation driven volume buffering effect has previously been described for simpler multicomponent condensates ^76^ and it could play a key role in stabilizing the environment of soluble proteins inside cells against external perturbations.

### Weak interactions are regulated when cells adapt to stress

Our comparison between exponentially growing and starved cells revealed that cells can modulate weak interactions when they adapt to new conditions. This complex proteome re-arrangement is likely a combination of global and protein specific effects. We observe a general tendency of soluble proteins to become more condensation-prone in starved cells while condensation-prone proteins become more soluble overall. These global shifts might be caused by a change in the chemical properties of the cytoplasm. In yeast, a drop in intracellular pH has been shown to drive the formation of reversible enzyme filaments in starvation conditions ^77–79^. While we adjust extract pH before performing PEG induced demixing experiments, it is possible that pH induced changes are not fully reversed in the time between pH adjustment and extract demixing, hence leading to the increased condensation propensity of normally high soluble proteins. Similarly, the increased solubility of condensation-prone proteins might be caused by altered solute properties of the cytoplasm. Starved yeast cells for example accumulate the storage carbohydrate trehalose ^80^, which has been shown to increase protein solubility and stress resistance ^81^.

Apart from such global changes, solubility of proteins could be regulated at the individual protein level by post translational modifications, chaperones or binding to specific metabolites. In addition, specific protein features might influence how proteins respond to altered conditions. Our analysis for example revealed that the presence of disordered regions makes proteins more likely to change solubility in either direction when cells enter starvation. This observation might be explained by the recent observation that disordered protein regions experience a disproportionate change in phosphorylation status when human cells adapt to growth at altered temperatures or osmolarity ^82^. How these different processes mechanistically modulate weak interactions as cells adapt to stress remains to be determined.

### Proteome demixing allows a differential cellular response to perturbations

Our approach does not measure protein activity, but protein condensation and the resulting changes in local environment are expected to affect protein function. Condensates have been shown to influence protein activity and act as biosensors for crowding, pH and temperature changes ^69,83,84^. Therefore, we expect that the function of condensation-prone proteins would be influenced by biophysical changes more than the function of proteins that remain soluble. We show that the core protein synthesis machinery and its key regulator TORC1 are condensation-prone, providing an attractive explanation for the long-standing observation that protein synthesis rates decline in overcrowded conditions ^29,57–59^. Whether the crowding dependent TORC1 condensates correspond to the previously described supramolecular TORC1 assemblies that ensure TORC1 inactivation during starvation remains to be determined ^85,86^.

While in the past studies have often focused on proteins that form condensates, our work also highlights the importance of high protein solubility. We demonstrate this for the stress induced MAP kinase Hog1, which needs to remain soluble to mount a rapid transcriptional response to hyperosmotic compression. Likewise, many metabolic enzymes and stress-induced proteins remain soluble at high crowding, suggesting that these processes are less affected by environmental fluctuations. In conclusion, our data show that weak interactions play a key role in organizing the intracellular space by promoting the formation of dynamic, environment dependent supramolecular assemblies. We suggest that this dynamic self-organization allows cells to couple some processes to external perturbations while others continue to work robustly, despite environmental fluctuations. When cells adapt to new conditions, these properties might get adjusted to ensure an optimal cellular response in the new environment.

### Limitations

In principle our fractionation method could allow identification of co-condensation networks of proteins. However, the current data quality does not allow this. Clustering algorithms tend to cluster proteins from known condensates close together, but often split in two or more sub-clusters, and usually interspersed with proteins from other modules. Therefore, we currently cannot say whether our data reflects the formation of many different condensates with specific composition or whether the proteome assembles into unspecific condensates with only weak enrichment of individual proteins. Similarly, our method cannot make any statements about the material properties of the identified condensates. The use of synthetic crowders carries the risk of introducing crowder-specific artefacts for some proteins. We tried to mitigate this risk by using orthogonal approaches and by comparing our data with other datasets.

## DATA, CODE, AND MATERIALS AVAILABILITY

The mass spectrometry proteomics data have been deposited to the ProteomeXchange Consortium via the PRIDE partner repositoryincluding analysis results in the format of .sne files and the related .tsv reports as used in the downstream data analysis.

Data associated with the mass spectrometry part of the manuscript, including specific versions of proteomes and databases used for the analysis, are stored in a Zenodo object.

Code used to analyze the data and generate figure panels in R and python is stored on Zenodo at or is available on the lab’s Github page.

All data, code and related objects will be made public upon publication of the manuscript.

## ACKNOWLEDGEMENTS

We thank Robbie Loewith, Eric Dufresne, Paolo Arosio for fruitful discussions. We are grateful to Antje Dittmann, Tobias Kockmann, the FGCZ and Katarzyna Makasewicz for technical support and suggestions. We thank Arthur Molines, Yves Barral, Madhav Jagannathan, Pedro Beltrao and Matthieu Piel for feedback on the manuscript and to Michele Marass for providing editorial advice during the publication process. GN was supported by a BIF PhD fellowship. FZ was supported by a MSCA global postdoctoral fellowship (101154995). SR was supported by the Max Planck Society and the Deutsche Forschungsgemeinschaft Project-ID 528483508 – FIP 12 (SR). GEN was supported by ETH Zurich and the Swiss National Science foundation by grants 310030_212660 and Eccellenza PCEFP3_187003.

## AUTHOR CONTRIBUTIONS

Conceptualization: GN, FZ, LW, PP, GEN; Methodology: GN, FZ, LW, LG, AAR, IK, TM, GEN; Investigation: GN, FZ, LW, LY, MB, AAR, LG; Visualization: GN, FZ, LW; Funding acquisition: GN, FZ, AAR, TM, SR, PP, GEN; Project administration: GEN; Supervision: SR, PP, GEN; Writing – original draft: GN, FZ, LW, GEN; Writing – review & editing: GN, FZ, LW, AAR, TM, SR, GEN.

## MATERIALS AND METHODS

### Yeast strains and culture conditions

*S. cerevisiae* strains are derivatives of W303 or BY4741 as indicated A list of yeast strains and strain backgrounds used in this study is provided in supplementary *Table S3.*Yeast strains were generated using PCR based transformation as described in ^89^. To generate the Hog1-GFP-LCD strain, the sequence coding for the LCD domain of Dbp1 ^90^ was cloned into the pFA6a-GFP(S65T)-kanMX6 (39292; Addgene) digested with AscI (R0558, New England Biolabs) and BstBI (R0519, New England Biolabs) using the Gibson Assembly technology (A46627; Invitrogen). The oligo used in the Gibson Assembly is listed in *Table S2*.

Cells were grown in standard conditions using either synthetic complete medium supplemented with 2% glucose (SCD) or yeast peptone medium supplemented with adenine (0.055 mg/mL) and tryptophan (0.8 mg/mL) and 2% glucose (YPD). Cells were grown in flasks at room temperature (live-cell imaging) or at 30C (extract preparation). Cell growth was monitored by measuring absorption at 600nm (OD_600_) using a spectrophotometer.

### Cell extract preparation

Yeast extracts were generated essentially as described in ^91^. In short, cells were inoculated overnight at 30 °C and diluted to OD_600_ < 0.1 in 1L fresh SCD medium in the morning. Cells were harvested at an OD_600_ between 0.5 and 0.7 by filtration using 0.8um-pore Mixed Cellulose Esters Membrane (Merk, AAWP09000). The filter cake was washed with 50 mL of a mixture of freshly prepared 100mM K_2_HPO_4_ in SCD. Subsequently, cells were scraped into a liquid-nitrogen filled 50 mL Falcon tube (Corning, 10788561) to flash-freeze them. HeLa cell pellets were purchased from Ipracell (CC-01-10-10). The frozen pellet was broken into smaller pieces before proceeding to cryo-milling.

Frozen cell pellets (yeast or HeLa) were processed for 1 cycle of 1 minute at a rate of 10 cycles per second in a cryogenic grinder (Spex 6870). The cell powder was transferred to Safe-Lock tubes (Eppendorf, 0030120086) and these tubes were kept in LN2 until all powder had been distributed to the tubes. The samples were gently thawed at 4 °C with shaking for 10-20 min before proceeding to extract clearance through centrifugation for 30 min at 16’000g at 4 °C in a refrigerated tabletop centrifuge. The supernatant was collected and pH adjusted to pH=7.1+-0.1 using 5M KOH. An additional centrifugation step of 5 min x16’000g at 4 °C was used to clear the samples from precipitate and floating lipid residues after pH adjustment. In all cases, protein concentration was determined by Bradford assay (Bio-Rad Protein Assay Dye Reagent Concentrate, #5000006) as described in ^92^. If proceeding to mix the lysate with PEG, an additional centrifugation step of 20 min x10’000g at 4 °C was included to improve data quality by removing everything that would pellet without PEG under the same conditions used after demixing induction.

### *X. laevis* growth and egg extract preparation

The Xenopus frogs used in this study are part of the Xenopus colony maintained at the Max Planck Institute for Infection Biology in Berlin, and were obtained from NASCO (Fort Atkinson, WI). Frogs were maintained in a recirculating tank system with monitored temperature and water quality at 18–20°C. All experimental protocols involving frogs were performed in accordance with national regulatory standards and ethical rules and reviewed and approved by the LaGeSo under Reg.-Nr. E 0051/25.

Low-speed metaphase-arrested egg extracts were prepared from Xenopus eggs arrested in metaphase of meiosis II as described previously ^93^. In brief, *X. laevis* frogs were primed with 100 U of pregnant mare serum gonadotrophin (PMSG) 3-7 days before the experiment and were boosted with 1000 U human chorionic gonadotrophin (HCG) to induce egg laying. Eggs arrested in the metaphase stage of meiosis II were collected, dejellied using L-Cysteine and fractionated via centrifugation. The cytoplasmic layer was isolated and supplemented with Cytochalasin (10 µg/mL final) and Complete EDTA-free protease inhibitor (1:100). Cycling to interphase was induced by the addition of CaCl2 to 0.6 mM and Cylcohexamide to 10 µg/mL final concentration.

### PEG-induced phase separation

PEG (35kDa, BioUltra, Sigma) dissolved in K-PBS (154 mM Kcl (Sigma), 2.2 mM KH2PO4 (Merck), 10 mM K2HPO4 (Merck), pH 7.1) at 5x the final intended concentration was pipetted in PCR tubes (Thermo Scientific, AB-1182 for the yeast experiment, Axygen maximum Recovery – Sigma-Aldrich AXYPCR02LC for the HeLa and Xenopus experiments). Lysate was added on top at 4x the volume of the PEG. Each sample was mixed by gentle but quick repeated pipetting and vortexing.

After 2 min equilibration at the chosen temperature, samples were placed in a temperature-controlled tabletop centrifuge and centrifuged at 10’000 rcf for 20 min. After that, samples were removed, and the supernatant was removed while measuring its volume by pipetting. We added a known volume of water to solubilize the pellet for ease of handling and measured its volume by pipetting.

After this step, we measured the protein concentrations in the supernatant and in the resuspended pellet solution by Bradford assay (see above). We use these concentrations and volumes to calculate the bulk protein partitioning into the pellet and dilute fraction and compare these numbers to non-demixed samples to make sure no protein was lost during demixing and that our volume and concentration measurements are accurate.

### Demixing reversibility experiment

Yeast extract demixing was induced as described above using a final PEG concentration of 5%. After that, the sample was diluted to a final PEG concentration of 0.5%. This was done with three distinct solutions: Potassium-based PBS (K-PBS), cell extract from wild type yeast, or with the flowthrough of the extract (FT) after it passed through an Amicon filter (Ultra-0.5, 10 kDa cutoff, UFC501096). Flowthrough and PBS dilute both PEG and proteins. The extract from WT cells ensures that concentrations of proteins and small molecules don’t change, with only PEG being diluted. Because we use WT extract (Dcp2 not fused to GFP) the average Dcp2-GFP signal is lower, but all the fluorescent signal comes from the previously condensed Dcp2-GFP. After dilution the solution appeared macroscopically clear and homogeneous in all conditions.

### Mass-spectrometry sample preparation

After PEG induced sample demixing, we prepare a set protein amount (unless otherwise specified, 200 µg) for digestion in 10 µL, bringing to volume with MilliQ water, in thin-walled PCR tubes (Thermo Scientific, AB-1182). Samples were diluted with 40 µL Sodium Deoxycholate (in the following: DOC, Fluka) in ammonium bicarbonate (in the following: AB, Sigma) to final concentrations of 5% DOC and 10 mM AB.

Samples were heat-treated on a thermocycler for 5 min at 99C to prevent sample degradation, then cooled for 5 min on ice. Samples were then transferred to either protein LoBind Eppendorf tubes (0030108442, 0030108450), or to plates (VWR, VWRI732-3806 and Axygen, P-2ML-SQ-C-S), and processed as follows: reduction with 5 mM dithiothreitol (in the following: DTT, Applichem) for 30 min at 37 °C, alkylation with 20 mM iodoacetamide (in the following: IAA, Merck) for 30 min at RT in the dark, both with shaking.

#### For all experiments except the stationary vs. exponentially-growing phase comparison

Samples were diluted in 10 mM AB and digested overnight with 1:66 Sequencing grade modified trypsin (Promega, V5113).

The next morning, samples were acidified with phosphoric acid (Merck) and filtered through 0.2 µM plates (Pall Acroprep Advance) to stop the reaction and remove the DOC.

From here on all mentioned solvents in this section are from Fisher, Optima LCMS grade. Buffer composition in this section: Buffer A: 0.1% FA in H2O, Buffer B: 0.1% FA in 50% ACN, 50% H2O. Buffer C: 5 mM KH2PO4, 25% ACN and pH adjusted to 2.8 with phosphoric acid. Buffer D: 5 mM KH2PO4, 300 mM KCl, 25% ACN and pH adjusted to 2.8 with phosphoric acid.

Samples underwent strong cation exchange on BioPureSPN 96-Well Midi Plates (The Nest group/Poly LC) to remove PEG on a vacuum manifold. Before loading the peptides, the plate was primed with the following washes: 200 µL Methanol, 400 µL H2O, 2×400 µL Buffer D, 2×400 µL Buffer C. After loading peptides, we performed 2×400 µL Buffer C washes followed by elution in a fresh plate with 400 µL Buffer D. Samples were dried in the speedvac and rehydrated in buffer A: 0.1% FA (Fisher, A117-50) in H2O.

#### For the stationary vs. exponentially-growing phase comparison

In this experiment, only 100 µg of protein per sample were processed. Reduction alkylation was performed as described above, then samples were purified from PEG and digested following the SP4 protocol as in ^94^, in the glass-bead version, using ACN as a precipitating agent. Digestion of each samples was performed in 100 µL of 100 mM AB containing 1.5 µg trypsin, for 18h at 37C. Recovery of the supernatant by centrifugation and an additional 100 µL washing of the beads with 100 mM AB was followed by acidification with FA.

#### For all experiments

After acidification, samples were desalted with Pierce Peptide Desalting Spin Columns (89851). We adapted the vendor protocol with the following washes: 1×300 µL Methanol, 2×300 µL Buffer B, 3×300 µL Buffer A. After sample loading, we washed 3×300 µL with Buffer A and eluted with 2×300 µL of buffer B into fresh 2 mL Protein LoBind tubes (Eppendorf, 0030108132). Steps before peptide loading were performed with 1 min x5000 rcf centrifugations at RT, peptide loading and after at 1 min x3000 rcf at RT. Samples were dried, then resuspended in Buffer A prior to acquisition.

#### Mass-spectrometry acquisition

After rehydration, samples were measured on a Lunatic or Nanodrop and around 100-150 ng peptides/sample were injected. LC-MS/MS analysis for yeast samples (except the stationary/exponentially-growing phase comparison) was performed on an Orbitrap Exploris 480 mass spectrometer (Thermo Scientific) coupled to an Acquity UPLC M-Class system (Waters). Peptides were separated using a C18 reversed phase nanoEase M/Z HSS T3 Column (100Å, 1.8 µm, 75 µm X 250 mm, Waters, 186008818), using a 120-minute linear gradient from 5% to 30% buffer B at a flow rate of 300 nL/min (buffer A: 0.1% [v/v] formic acid, buffer B: 0.1% [v/v] formic acid in acetonitrile). The mass spectrometer was operated in data-independent acquisition (DIA) mode with the following parameters: one MS1 scan (350-1150 m/z, 60000 resolution, 200% normalised AGC target, 100 ms maximum injection time), followed by 41 variable MS2 windows from 350 to 1150 m/z with 1 m/z overlap (30000 resolution, 200% normalised AGC target, 54 ms maximum injection time). Ions were fragmented with HCD (NCE 30%).

LC-MS/MS analysis for human and frog samples was performed on an Orbitrap Exploris 480 mass spectrometer (Thermo Scientific) coupled to a Vanquish Neo UHPLC system (Thermo Scientific). Peptides were separated using a C18 reversed phase column (75 μm x 400 mm (CoAnn), packed in-house with ReproSil Gold 120 C18, 1.9 μm (Dr. Maisch GmbH)), using a 120-minute non-linear gradient from 1% to 50% buffer B at a flow rate of 300 nL/min (buffer A: 0.1% [v/v] formic acid, buffer B: 0.1% [v/v] formic acid, 80% [v/v] acetonitrile). The mass spectrometer was operated in data-independent acquisition (DIA) mode with the following parameters: one MS1 scan (330-1650 m/z, 120000 resolution, 300% normalised AGC target, 20 ms maximum injection time), followed by 50 variable MS2 windows from 330 to 1650 m/z with 1 m/z overlap (30000 resolution, 2000% normalised AGC target, 64 ms maximum injection time). Ions were fragmented with HCD (NCE 27%).

LC-MS/MS analysis for the stationary/exponentially-growing phase yeast experiment was performed on an Orbitrap Exploris 480 mass spectrometer (Thermo Scientific) coupled to a Vanquish Neo UHPLC system (Thermo Scientific). Peptides were separated using a C18 reversed phase column (75 μm x 400 mm (CoAnn), packed in-house with ReproSil Gold 120 C18, 1.9 μm (Dr. Maisch GmbH)), using a 60-minute non-linear gradient from 1% to 43.7% buffer B at a flow rate of 300 nL/min (buffer A: 0.1% [v/v] formic acid, buffer B: 0.1% [v/v] formic acid, 80% [v/v] acetonitrile). The mass spectrometer was operated in data-independent acquisition (DIA) mode with the following parameters: one MS1 scan (330-1650 m/z, 120000 resolution, 300% normalised AGC target, 20 ms maximum injection time), followed by 50 variable MS2 windows from 330 to 1650 m/z with 1 m/z overlap (30000 resolution, 2000% normalised AGC target, 64 ms maximum injection time). Ions were fragmented with HCD (NCE 27%).

#### Data analysis and post-analysis

All data was analyzed with Spectronaut (Biognosys) using standard parameters in DirectDIA mode. We validated protein integrity by running semi-specific searches on a reduced number of samples in each experiment (data not included).

Analysis was done using the UniProt reference proteome for all experiments. For Xenopus experiments, we performed an additional analysis using the Xenbase ^95^ v9.2 assembly of the proteome, for easier comparison to the data included in ^96^. All proteomes are available in the PRIDE repository associated with this paper.

The post analysis was done in R (v4.3.2) in Rstudio (2023.12.1.402), in a dedicated project and using Git for version control and renv (v1.1.5) to manage the environment. All data processing has been done through R. The main packages used for the MS-specific analysis are Protti ^97^ for reading peptide-level reports from Spectronaut, to normalize data and to calculate protein abundances. A minimal package called protRatio was developed in house to facilitate calculation of supernatant to total calculations. It is available as source package and code at Github. All data analysis notebooks are stably available on Zenodo including the protRatio version used in the paper. Please see the materials availability statement for the respective DOIs.

For the analysis pipeline, we used precursor-level raw ms2 intensities from Spectronaut to calculate a total per-sample intensity. We then filtered out precursors that are detected sparsely or at very low intensity, and those common to multiple proteins groups (the minimal set of proteins that explains the observed peptide evidence, where individual proteins cannot be distinguished). The remaining intensities were used to calculate protein group specific abundances using the Protti package ^97^. These abundances were normalized by the previously calculated total run intensities and multiplied with the per-condition total protein mass estimate, that we measured previously as described above, to obtain an absolute estimate of protein mass abundance (only of the protein group-specific peptides). Abundance estimates from paired supernatant and pellet samples were used to calculate the percent of each protein group in the dilute fraction, and the average of the fraction at each PEG concentration was used to calculate the Protein Solubility Score by trapezoidal integration (area under the curve or AUC). Total protein abundance estimate variability was used to exclude proteins where total abundances (supernant plus pellet) were inconsistent. To avoid inflation of the solubility scores in case of cross-run contamination, we force the ratio to 0 if only 1 out of 4 replicates has non-zero ratio at a given PEG concentration. We also calculate confidence intervals for the integral of the curve through bootstrapping (reported in data S2).

Data was further compared to publicly available databases such as Uniprot ^98^, PhaSepDB ^99^, PhaSePred^25^, ComplexPortal ^100^, the Alliance of Genome Resources ^101^, MobiDB ^102^, Alphafold ^103^, and cited papers as described in the text and figure legends. In particular: membrane association and subcellular localization were taken from Uniprot, protein net charge at neutral pH was calculated using the Peptides package in R ^104^, protein structural composition was taken both from ^68^ or calculated with a custom script based on the Protti package combined with AlphaFold scores (not reported in figures, but available in additional scripts). GSEA was conducted using the fgsea package in R, using a wrapper function modified from the Github repo associated to ^105^, using a customized gene set included in the data repository associated with this paper ^87^.

Differential abundance calculations were performed in R using the protti package ^97^. Protein abundances used for differential abundance calculations were obtained summing all precursor-level intensities for protein-group specific peptides that passed our quality control, after normalization, all through package protti. Differential abundance calculations on solubility (% soluble fraction of a protein across two conditions) where performed in the same way, setting an adjusted p-value threshold of 0.05 and a difference of at least 10 in solubility.

#### Yeast Live-Cell imaging and analysis

Yeast cells were inoculated overnight in SCD medium and refreshed at OD_600_=0.1. 200µL of cells were collected at OD_600_ =0.2-0.4 and loaded into Concanavalin A-coated (2mg/ml) Ibidi chambers (80806; Ibidi GMBH). Images were taken before and after the addition of 1M Sorbitol. Widefield pictures (Figure 2G; Figure 6E; Fig S4) were taken on a Nikon Eclipse Ti2 and an ORCA-fusion BT digital camera C15440, using the software NIS-Elements AR v5.42.03. Confocal pictures (Figure 1E; Figure 7E) were taken on a Nikon Eclipse Ti2 with the CSU-W1SoRa confocal scanner unit (Yokogawa) and an ORCA-fusion BT digital camera C15440, using the software NIS-Elements AR v5.42.03. z-stacks were acquired to cover the entire thickness of the cell. The number of foci per cell was quantified manually using FIJI. Cell-ACDC software (v1.4.31) ^106^ was used to quantify Hog1 nuclear migration, for performing cell-segmentation with YeaZv2 ^107^ and nuclear segmentation with pomBseen_nuclear ^108^.

#### Yeast cell extract imaging and analysis

Yeast cell extracts were prepared as described above. To modulate extract concentration, samples were concentrated up to 200 g/L by centrifugation at 14,000 × g for 30 min using Amicon centrifugal filters (Ultra-0.5, 10 kDa cutoff, UFC501096). The flow-through from the filtration was then used to dilute the sample to final concentrations as low as 5 g/L. In the case of PEG-induced phase separation, PEG and lysate were mixed as described above without equilibration time. The extract was mounted on a coverslip (CLS294875X25; Corning) and sealed with nail polish. Confocal pictures (Figure 1F; Figure 6C; Figure 7A; Figure S1A) were taken on a Nikon Eclipse Ti2 with the CSU-W1SoRa confocal scanner unit (Yokogawa) and an ORCA-fusion BT digital camera C15440, using the software NIS-Elements AR v5.42.03. FIJI was used to calculate the number of foci per field of view and the fluorescence intensity in the dense/soluble phases. Microscopy images were acquired on a confocal microscope as described in the yeast live-cell imaging section.

#### FCS

FCS measurements were performed on an inverted confocal fluorescence microscope (Leica SP8 STED, Leica Application Suite X [LAS X] software, version 1.0) using an HC PL APO CS2 63× 1.2 NA water immersion objective with a software-controlled correction collar (Leica, Wetzlar, Germany) and a hybrid detector optimized for single-molecule detection (HyD SMD). The confocal volume was routinely calibrated using Atto 488 free acid (D=400 um2/s). Typical calibrations yielded an effective volume of approximately 0.4 fL and a structural parameter around 8. Samples were excited with a 488 nm laser (white Light Laser, 80 MHz), and fluorescence emission was collected between 500 and 550 nm. Leica Application Suite X [LAS X] software was used to fit the correlation curves with a 1-component anomalous diffusion model to extract the diffusion coefficients. The diffusion coefficients measured at different overall protein concentration were fitted using with our FCS model to extract the parameters *D_0_, K^a^_eff_* and *κ*. *D_0_, K^a^_eff_* are fitted for each protein individually, and *κ* is fitted globally (see supplementary text).

#### Immunoblot analysis

Yeast cells were inoculated overnight in YPD medium and refreshed at OD_600_=0.1. Once at OD_600_=0.4, cells were treated with Sorbitol 1M (final concentration in YPD). At the indicated time points, 5mL-10 mL of cells was fixed with 5% trichloroacetic acid (TCA) for 10 minutes. TCA was removed by centrifugation; the pellet was washed with acetone and air-dried. Cells were lysed in 50 mM Tris-HCl, pH 7.5, 1 mM EDTA, 1 mM p-nitrophenyl phosphate, 50 mM dithiothreitol, 1 mM phenyl-methylsulphonyl fluoride, and 2 µg/mL pepstatin with glass beads in a fast prep. Samples were mixed with 1/3 volume of 3X sample buffer (187.5 mM Tris oh6.8, 6% β-mercaptoethanol, 30% glycerol, 9% SDS, 0.05% bromophenol blue) and boiled for 10 minutes before running NuPAGE Tris-Acetate gels (Invitrogen). For western blot analysis, proteins were transferred onto nitrocellulose membrane. The membrane was blocked with PBS (pH 7.4), 0.1% Tween-20, 3% Milk or 3% BSA before applying the primary antibodies. Antibodies used were: mouse monoclonal anti-Pgk1 (22C5D8, used at 1:1,000 dilution, Abcam) to detect Pgk1; mouse anti-Sch9-S758P (used at 1:25,000 dilution) and rabbit anti-Sch9 (used at 1:5,000) to detect respectively phosphorylated and total Sch9 were a kind gift of Robbie Loewith ^109^ (Department of molecular and cellular biology, Université de Genève); rabbit anti-p38-T180P/T182P (3D7, used at 1:1,000 dilution, Cell Signaling Technology) to detect phosphorylated Hog1; mouse anti-Hog1 (D-3, used at 1:1,000 dilution, Santa Cruz Technology) to detect total Hog1. The secondary antibodies used were: donkey anti-mouse IgG-HRP linked (NA931, used at 1:3,000 dilution, Amersham); donkey anti-rabbit IgG-HRP linked (NA934, used at 1:3,000 dilution, Amersham).

#### qPCR

5 OD_600_ units of cells were collected and frozen in liquid nitrogen. Pellets were re-suspended in 400 μL TES buffer (10 mM Tris pH7.5, 10 mM EDTA, 0.5%SDS) and 400 μL acid phenol and incubated for 45 minutes on a thermoshaker at 65 °C after addition of 100µL glass beads. The aqueous phase was separated from the phenol by centrifugation, carefully isolated and mixed with phenol for a second round of phenol extraction, followed by a chloroform extraction. RNA was cleared using the RNeasy MinElute Cleanup Kit (74204, Qiagen) with DNAse treatment (79254, Qiagen) according to the manufacturer’s instructions. cDNA was generated using 1,000 ng of RNA as a template for the Reverse Transcriptase reaction (18064014, Invitrogen). Gene expression was measured using the Kapa SYBR fast for LightCycler 480 (KK4611, Roche) and primer sets specific for each gene of interest. The qPCR primers used in this study are listed in *Table S2*.

#### NMR

We induced demixing of 40 µL PEG 25% with 160 µL of yeast lysate at 4C. After fractionation, pellets were resuspended with 50 µL of water, and volumes and protein concentrations were measured as described above. NMR samples were prepared by mixing either 10 µL of the 25% PEG solution, 10 µL supernatant or 60 µL resuspended pellet with 600 µL D2O, and 5 µL DMF. After transferring the samples to NMR tubes (DWK Duran economic 300 MHz tubes, 23-170-01-17), ^1^H NMR spectra were acquired using a 300 MHz instrument (300 Ultrashield, Bruker) at 298 K and averaged 16 times. The resulting data were analyzed using the MestreNova software (Mestrelab Research). The PEG concentrations were determined from the relative height of the PEG peak to the two DMF peaks and the known DMF concentration in the sample.

