## Supplementary text for "Weak interactions drive selective proteome demixing and tune the differential response to environmental perturbations"

### Derivation of the effective diffusion coefficient and bound fraction in a multi-partner cytoplasm

*1. System definition and hypotheses*

We consider a protein $X$ diffusing in the cytoplasm and that can interact with cytoplasmic partners $Y_i$, indexed by $i=1,\dots,N$. Each interaction is described by the reversible reaction

$$X + Y_i \underset{k_{off}}{\overset{k_{on}}{\rightleftharpoons}} XY_i$$

FCS measurements of protein $X$ in yeast extracts lead to autocorrelation curves that are well described by a single diffusive component from 0 to 80 g/L overall protein concentration (Fig. 2). This indicates that the exchange between free and bound states does not give rise to resolvable multiple diffusive populations. Therefore, the measured dynamics can be characterized by a population-averaged effective diffusion coefficient $D_{eff}$, which reflects both the cytoplasmic environment and the molecular interactions.

*2. Mass conservation and bound fraction*

Let $[X]$ be the concentration of free protein and $[XY_i]$ the concentration of protein bound to partner $Y_i$ ($i=1,\dots,N$). The total concentration of the protein is

**Equation 1**

$$[X]_{tot} = [X] + \sum_{i=1}^{N} [XY_i].$$

At equilibrium each complex satisfies

**Equation 2**

$$[XY_i] = K_i^a [X][Y_i] (i=1,\dots,N),$$

so that substituting Equation 2 into Equation 1 gives

$$[X]_{tot} = [X]\left(1 + \sum_{i=1}^{N} K_i^a [Y_i]\right).$$

We define the microscopic bound fraction for species $i$ as

**Equation 3**

$$f_i \equiv \frac{[XY_i]}{[X]_{tot}} = \frac{K_i^a [Y_i]}{1 + \sum_{j=1}^{N} K_j^a [Y_j]}.$$

The total bound fraction is therefore

**Equation 4**

$$f_b = \sum_{i=1}^{N} f_i = \frac{\sum_{i=1}^{N} K_i^a [Y_i]}{1 + \sum_{j=1}^{N} K_j^a [Y_j]}.$$

3. Effect of multiple bound states on the effective diffusion coefficient

Let $D_{free}$ denote the diffusion coefficient for free $X$ and $D_{bound,i}$ the diffusion coefficient of $X$ when bound to $Y_i$. The population-averaged effective diffusion coefficient (accounting for free and all bound states) is the weighted sum

**Equation 5**

$$D_{eff}=\left(1-\sum_{i=1}^{N} f_i\right)D_{free}+\sum_{i=1}^{N} f_i\,D_{bound,i}.$$

Substituting Equation 3 into Equation 5 yields

**Equation 6**

$$D_{eff}\&=\left(1-\frac{\sum_{j=1}^{N}K_j^a\,[Y_j]}{1+\sum_{j=1}^{N}K_j^a\,[Y_j]}\right)D_{free}+\sum_{i=1}^{N}\frac{K_i^a[Y_i]}{1+\sum_{j=1}^{N}K_j^a\,[Y_j]}D_{bound,i}$$

$$\&=\frac{1}{1+\sum_{j=1}^{N}K_j^a\,[Y_j]}D_{free}+\frac{1}{1+\sum_{j=1}^{N}K_j^a\,[Y_j]}\sum_{i=1}^{N}K_i^a\,[Y_i]D_{bound,i}$$

$$\&=\frac{D_{free}+\sum_{i=1}^{N}K_i^a\,[Y_i]D_{bound,i}}{1+\sum_{j=1}^{N}K_j^a\,[Y_j]}.$$

4. Expression in terms of an effective association parameter

We now introduce the total cytoplasmic protein concentration and the effective association parameter. The overall protein concentration in the cytoplasm, $P$, includes both the protein of interest and its interaction partners.

**Equation 7**

$$P\equiv\sum_{i=1}^{N}[Y_i]+[X],\,K_{eff}^a\equiv\frac{\sum_{i=1}^{N}K_i^a\,[Y_i]}{P},$$

We also define the affinity-weighted average bound diffusion coefficient

**Equation 8**

$$D_{bound,eff}\equiv\frac{\sum_{i=1}^{N}K_i^a\,[Y_i]D_{bound,i}}{\sum_{i=1}^{N}K_i^a\,[Y_i]}=\frac{\sum_{i=1}^{N}K_i^a\,[Y_i]D_{bound,i}}{K_{eff}^aP},$$

where $D_{bound,i}$ denotes the dilute-solution diffusion coefficient of the protein bound with $Y_i$.

Using Equation 5 and Equation 8, we can re-write:

**Equation 9**

$$D_{eff}=\frac{D_{free}+(K_{eff}^aP)D_{bound,eff}}{1+K_{eff}^aP}.$$

5. Incorporation of cytoplasmic viscosity

The effect of global cytoplasmic crowding on protein diffusion is complex, and several empirical expressions have been proposed in the literature. To avoid adopting a model without clear justification, we use a minimal phenomenological expression in which viscosity increases linearly with total protein concentration:

**Equation 10**

$$\eta(P)=\eta_0+\eta_1 P=\eta_0\left(1+\frac{\eta_1}{\eta_0}P\right)=\eta_0(1+\kappa P).$$

Using the Einstein relation, which states that diffusion is inversely proportional to viscosity, we obtain:

**Equation 11**

$$D_{free}(P)=\frac{D_{free}^0}{1+\kappa P}, D_{bound,eff}(P)=\frac{D_{bound,eff}^0}{1+\kappa P},$$

where $D_{free}^0$ and $D_{bound,eff}^0$ denote the corresponding diffusion coefficients in dilute solution.

The full expression for the effective diffusion coefficient measured by FCS is therefore

**Equation 12**

$$D_{eff}(P)=\frac{D_{free}^0+K_{eff}^a P D_{bound,eff}^0}{(1+K_{eff}^a P)(1+\kappa P)}.$$

### 6. Effect of transient interactions on the measured diffusion coefficient

In principle, the formation of a complex could introduce an additional state with diffusion coefficient $D_{bound}$, resulting in the full expression described above. In our experiments however, the autocorrelation curves are accurately described by a single diffusive component across all protein concentrations under 80 g/L. This indicates that FCS does not resolve the bound state as a separate species; instead, it detects a population-averaged diffusion coefficient. This might be explained by the complex environment of the cytoplasm, where proteins might form short-lived assemblies with a heterogeneous variety of partners, of different sizes and mobilities.

The affinity-weighted average bound diffusion coefficient therefore represents an average over many states that cannot be determined experimentally. It would therefore need to be added as a fitting parameter in the model. However, introducing $D_{bound,eff}^0$ leads to over-parameterization and fitting yields parameters that are not feasible. In contrast the free diffusion coefficient can be experimentally determined in very dilute conditions.

To prevent this over-parametrization, we consider an extreme case where binding of a protein to a cytoplasmic particle leads to complete immobilization of the protein. This would for example be the case if a protein binds to a large complex, mRNA, membrane or supramolecular assembly. Formally this would mean that $K_{eff}^a P D_{bound,eff}^0 \ll D_{free}^0$.

In this boundary condition, the effect of transient binding is captured entirely through the reduction of the free fraction. In this case, the effective diffusion coefficient can be approximated with:

**Equation 13**

$$D_{eff}(P) \approx \frac{D^0_{free}}{\left(1+K^a_{eff}P\right)(1+\kappa P)}.$$

This simplification allows us to determine the viscous coefficient through a global fit to the data, and determine the effective binding affinities between proteins and the cytoplasmic environment for each protein, shown in Figure 2.

Because proteins bound to other cellular components are still able to move to some degree, neglecting $K^a_{eff}PD^0_{bound,eff}$ introduces a systematic error. If we consider that $D^0_{bound,eff} > 0$, the binding affinity $K^a_{eff}$ would need to be larger than what we estimated when we neglected this term to explain the experimental data. In other words, when the bound protein fraction is not completely immobilized, more protein needs to be in the bound state to explain the experimental data. This means that the binding affinities determined using this approximation represent a minimal binding strength that are likely slightly below the real value.
