## Supplementary figures for "Weak interactions drive selective proteome demixing and tune the differential response to environmental perturbations"

**Figure S1.**

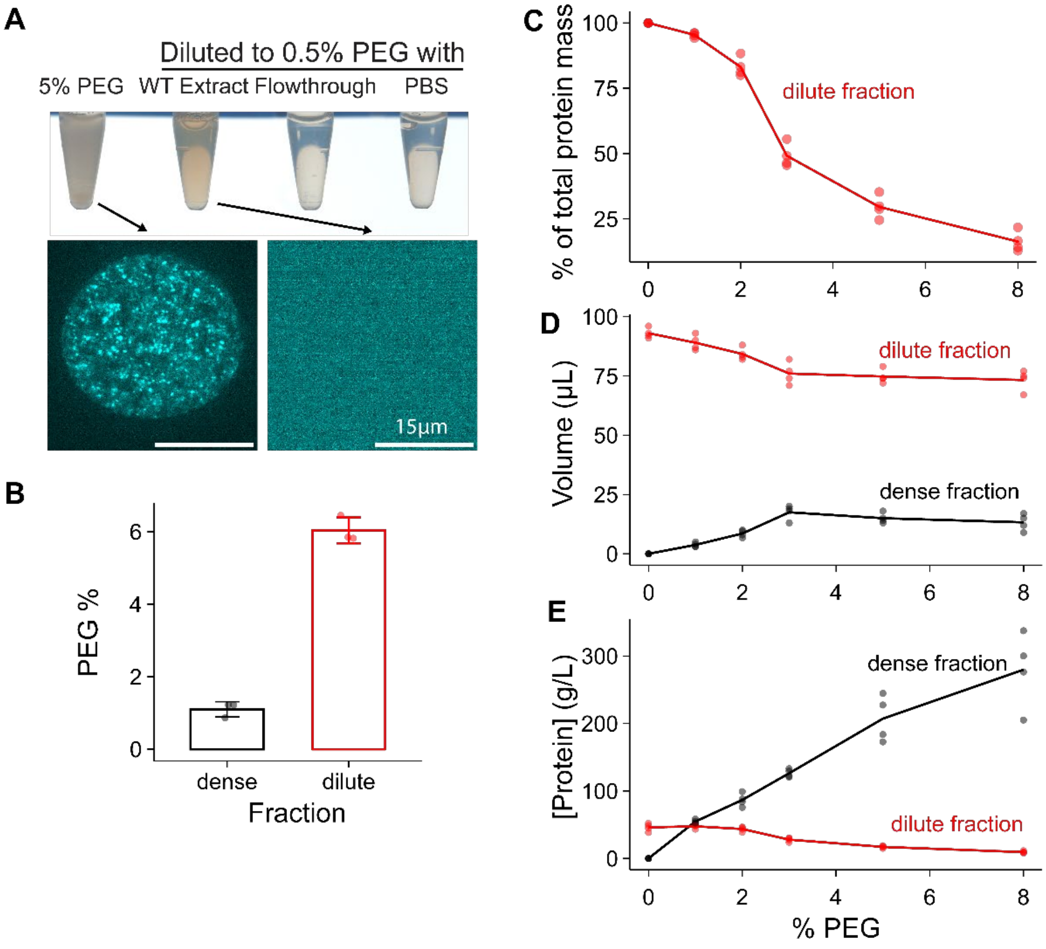


**Characterization of PEG-induced lysate demixing.**

**(A)** Images of yeast extract samples generated from the Dcp2-GFP strain after (5%) PEG addition, and after subsequent dilution to 0.5% PEG. Dilution was performed with 3 different solutions: cell extract, lysate flowthrough generated through 10 kDa MWCO filtration units, or PBS. All 3 methods yielded clear solutions. Microscopy images are shown at different contrast to highlight the difference in ultrastructure: both the large bubbles and the microscopic foci dissolve. **(B)** NMR-based measurements of PEG concentrations in the dense and dilute fractions, after inducing demixing of lysate with a total final concentration of 5% PEG at a temperature of 4 °C. **(C)** Quantification of the fraction of total protein mass remaining in the dilute fraction after PEG-induced demixing at different PEG concentrations. Total protein mass was quantified by measuring the volume (D) and the protein concentration (E) of the two fractions independently. **(D)** Quantification of the volume of the dense and dilute fractions after PEG-induced demixing at various PEG concentrations with a micropipette. Total expected volume 100 µL (80 of lysate and 20 of PEG stock solution). **(E)** Quantification of the protein concentration in the dense and dilute fractions after PEG-induced demixing at various PEG concentrations. Four independent measurements are shown.

**Figure S2.**

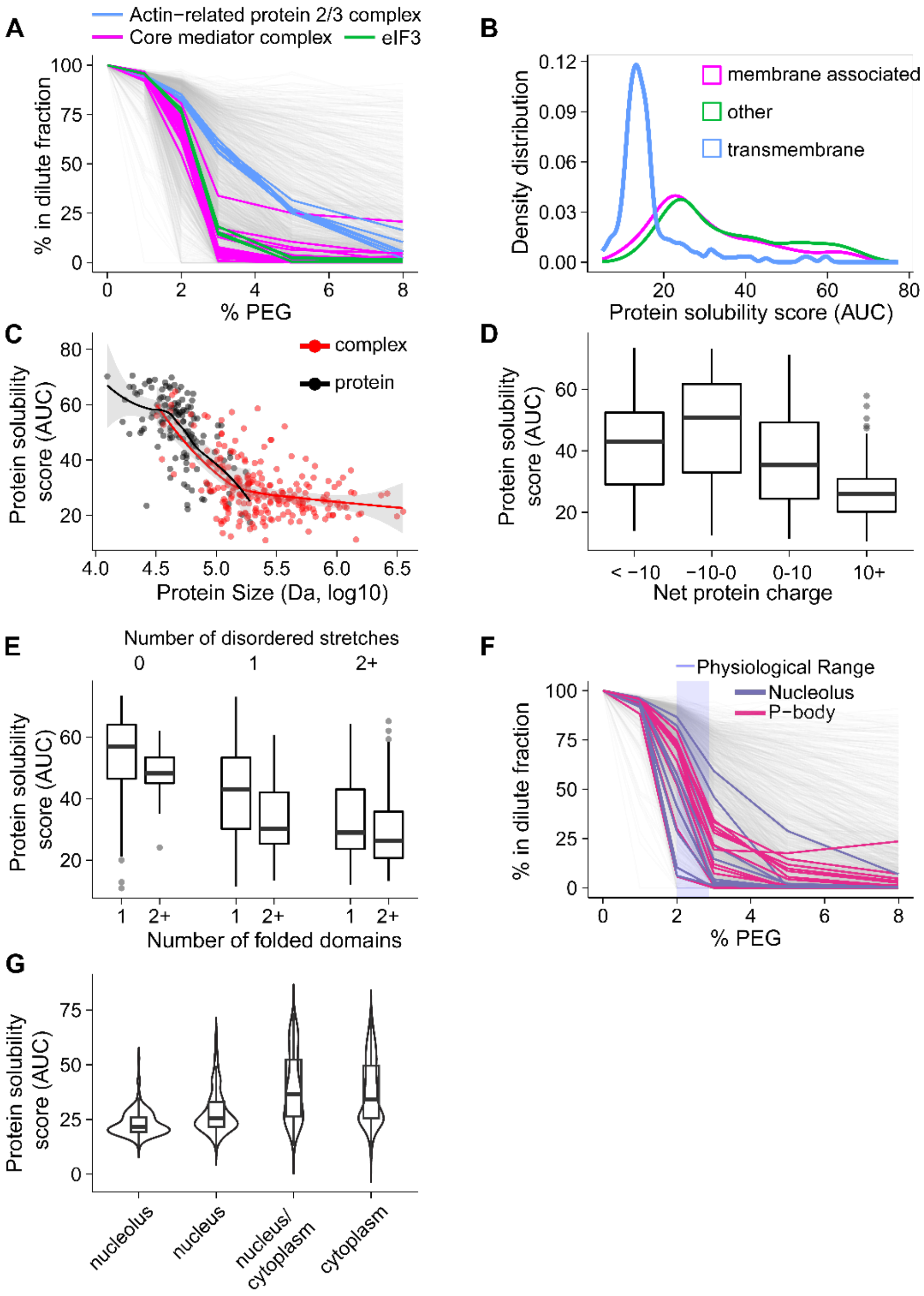


**Protein features correlated with condensation propensity.**

**(A)** Same data as shown in fig. 1B, but proteins that are part of the indicated complexes (Complex Portal [1]) are individually highlighted. **(B)** Distribution of protein solubility scores for proteins depending on their annotated relationship to membranes. Membrane-associated proteins contain all proteins having gene ontology terms containing “membrane” according to the UniProt [2] gene ontology annotation. **(C)** Protein solubility score plotted as a function of protein or complex (if applicable) size. For individual proteins (black), only proteins annotated as monomers or homomers, with no other annotated interactions (Uniprot [2]) are considered in this case. Complex mass is estimated from ComplexPortal-reported [1] composition and stoichiometry, setting the stoichiometry at 1 if unknown. **(D)** Distribution of protein solubility scores, for proteins that are not reported as being part of heteromeric complexes (Uniprot [2]), binned by net charge estimated at pH 7. **(E)** Distribution of protein solubility scores, for proteins that are not reported as being part of heteromeric complexes (Uniprot [2]), binned by the number of disordered stretches and folded domains identified by Romero-Perez *et al.* [3] through the DODO algorithm (qualitatively identical results obtained with the dataset with Chainsaw algorithm, and with alternative, in-house methods of determining folded domains and disordered stretches). **(F)** Same data as shown in fig. 1B, showing mass fraction of each protein remaining in the dilute fraction at the indicated PEG concentration. Proteins reported to associate with the indicated membrane-less organelles (PhaSepDB [4]) are individually highlighted. **(G)** Distribution of yeast protein solubility scores by intracellular localization.

**Figure S3.**

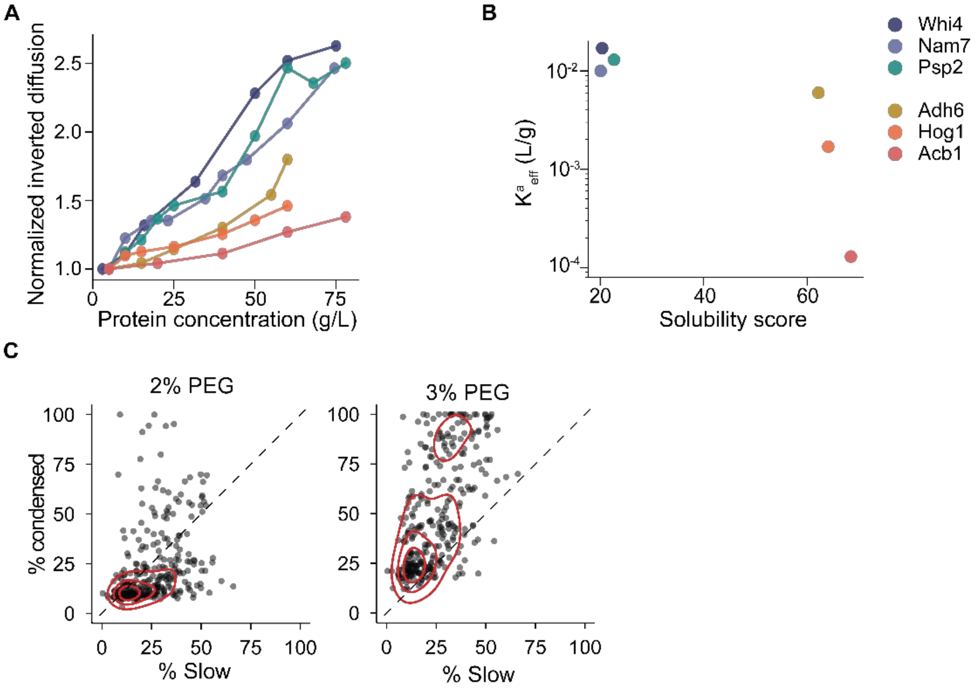


**FCS diffusion modeling.**

**(A)** Inverted diffusion coefficients of proteins ($1/D_{eff}$) normalized to the lowest concentration to remove protein-specific geometric contributions, as a function of protein concentration. This reports on the scaling of the protein-specific apparent viscosity through Stokes-Einstein, which does not consider protein interactions. The fact that this value is differently affected by extract concentration for different proteins shows that viscosity alone cannot explain the diffusion behavior of these proteins in concentrated solutions. **(B)** Effective binding equilibrium coefficients extrapolated from fitting of the data in Fig. 2A for each protein expressed in units of L/g. **(C)** Comparison of the fraction of protein partitioning to the condensed fraction in yeast extract at the indicated PEG concentration plotted against the slow moving fraction for each protein determined by in-cell FCS by Fukuda *et al.* [5].

**Figure S4.**

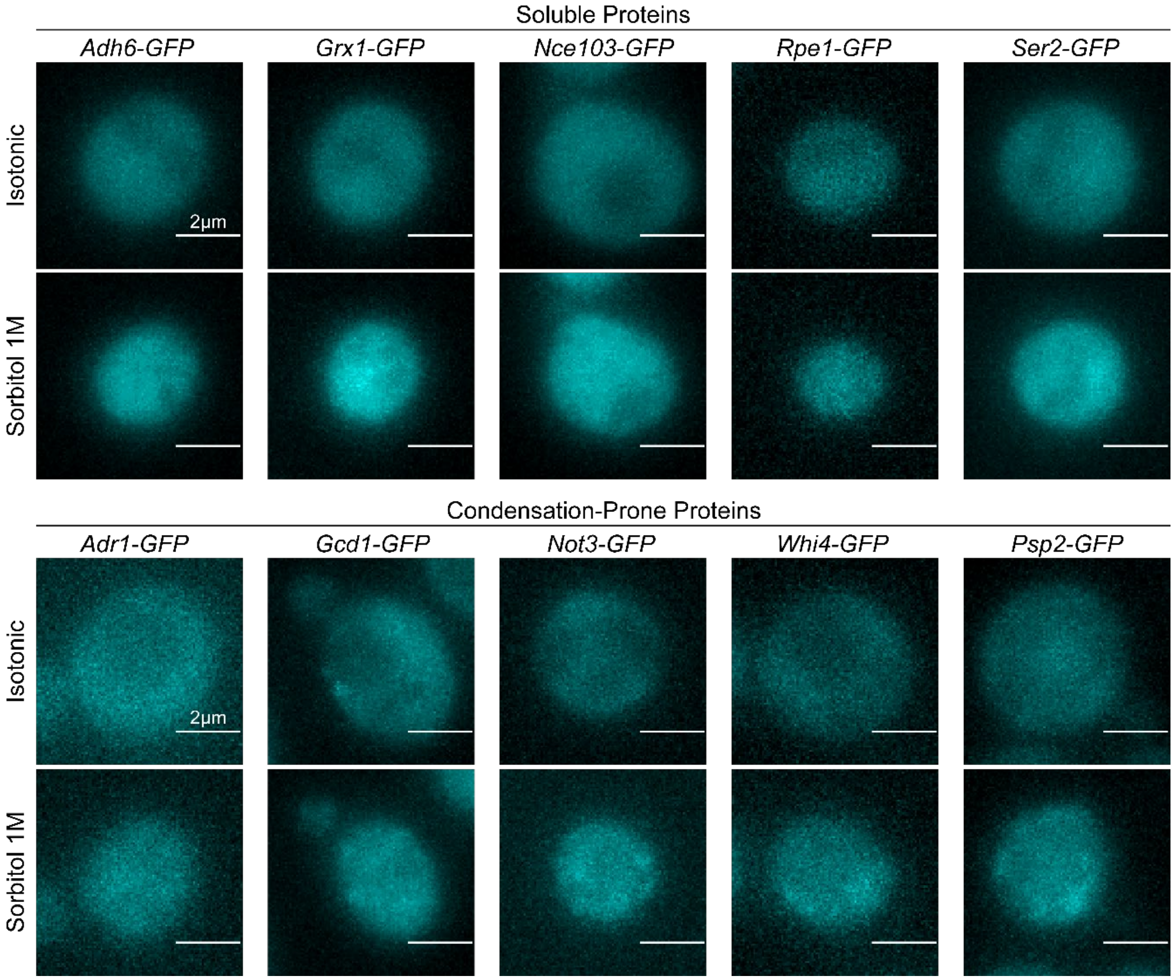


**Hypertonic stress and protein organization in cells.** Additional confocal images of yeast cells expressing GFP-tagged proteins with low or high solubility score (AUC) before and after 30 minutes from Sorbitol addition. The quantification of the phenotype is shown in Fig. 3F.

**Figure S5.**

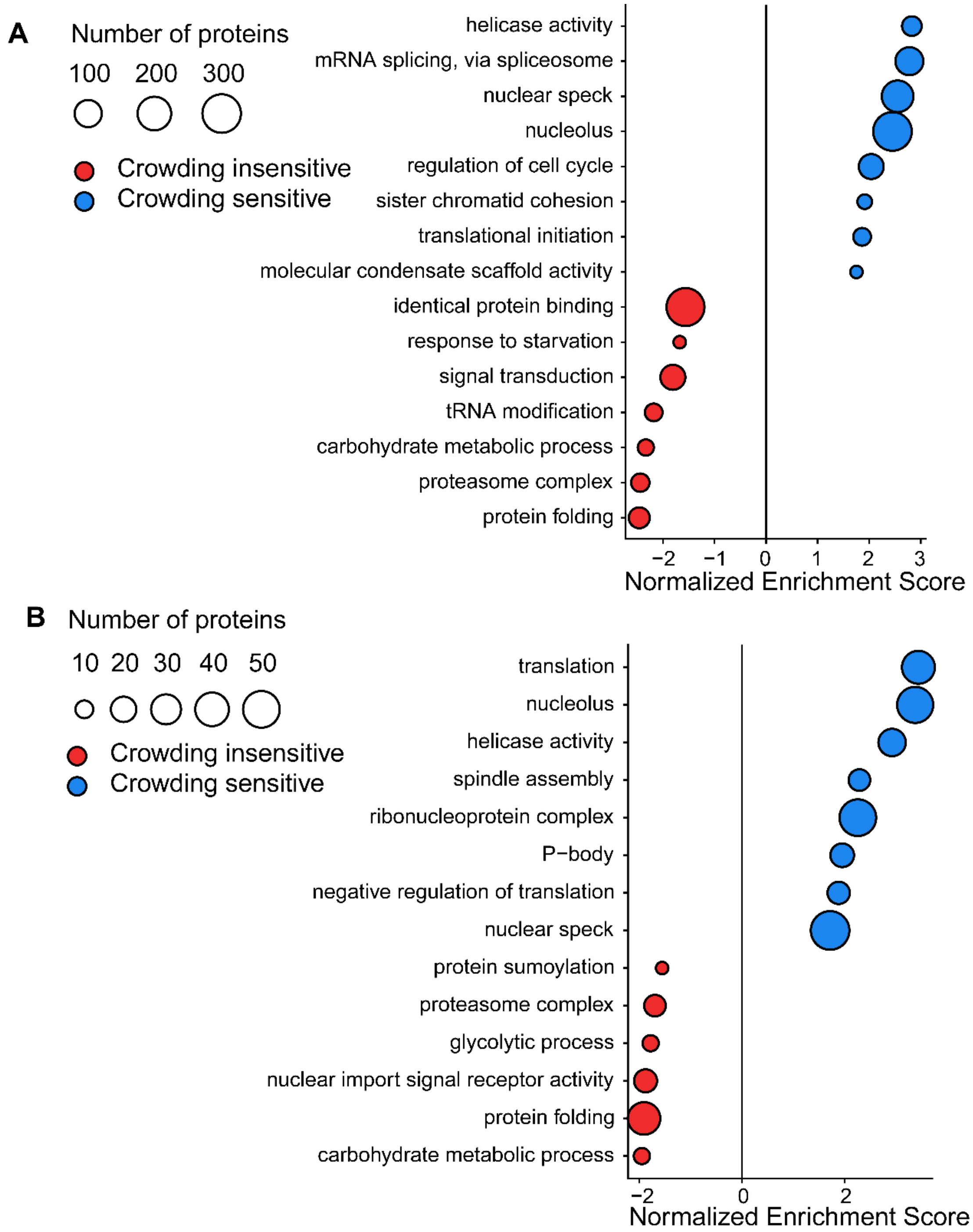


**Gene set enrichment analysis in other species.** Selected results of the single sample GSEA projection analysis on a ranked list of protein solubility in **(A)** HeLa cells and **(B)** *X. laevis* egg extract. In the xenopus case, only one gene per protein group (arbitrarily chosen) was included in the analysis to avoid extensive double-counting due to tetraploidy, which would strongly bias the GSEA results.

**Figure S6.**

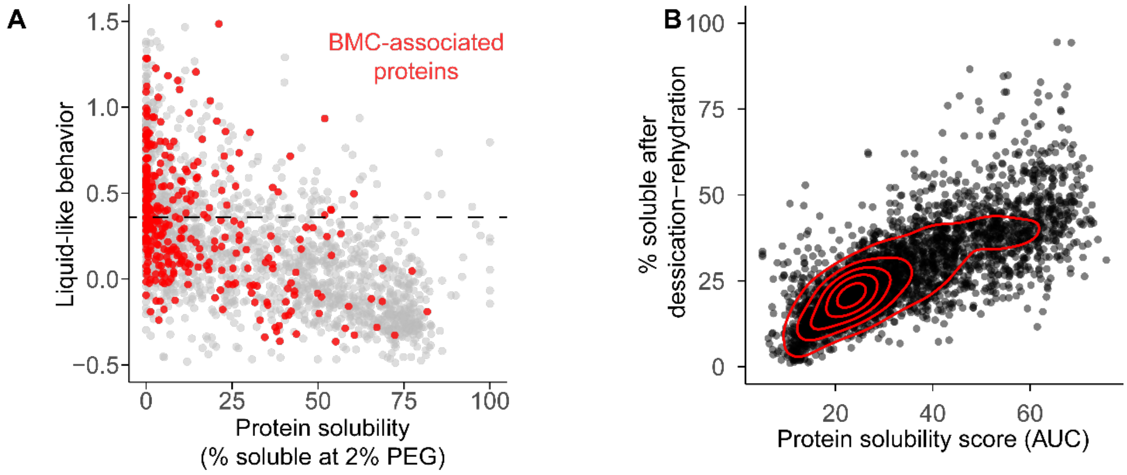


**Comparison with other proteome-wide studies.**
**(A)** Comparison between protein solubility in Xenopus egg extract in the presence of 2% PEG and the liquid-like behavior score from Keber *et al.* [6]. The horizontal dashed line corresponds to their threshold for high-confidence liquid-like mesoscale assembly components. **(B)** Comparison of our protein solubility score, obtained through a reversible, PEG-induced condensation profiling, with the irreversible aggregation following a complete desiccation-rehydration cycle performed on similar yeast extracts obtained by Romero-Perez *et al.* [3].

**Table S1. (separate file)**

Full protein partitioning and solubility score data.

**Table S2. (separate file)**

GSEA analysis results (only up to adjusted p-value of 0.1) for yeast, frog and HeLa protein solubilities.

**Table S3.**

| Figure panels | Genotype | Background | Source | Strain number |
|---|---|---|---|---|
| 1B-F, 1I-J, 3A, 4A, 4G | MATa, ade2-1, leu2-3,112, ura3, trp1-1, his3-11,15, can1-100, GAL [psi+] (**W303 WT**) | W303 | Neurohr lab | yGN611 |
| | MATa, his3Δ1, leu2Δ0, met15Δ0, ura3Δ0 (**BY4741 WT**) | BY4741 | Gift from Yves Barral | yGN3021 |
| 1G-H, S1A | MATa, Dcp2-mEGFP::caURA3 | W303 | Gift from Karsten Weis | yGN3330 |
| 1G-H | MATa, edc3Δ::LEU2, pat1Δ::KANmx, Dcp2-mEGFP::caURA3 | W303 | Neurohr lab | yGN3352 |

| | | | | |
|---|---|---|---|---|
| 3E-F | MATa, GCD1-GFP::HIS | BY4741 | Yeast GFP CC | yGN3376 |
| 3E-F | MATa, NOT3-GFP::HIS | BY4741 | Yeast GFP CC | yGN3377 |
| 2A, 2C-E, 3E-F | MATa, WHI4-GFP::HIS | BY4741 | Yeast GFP CC | yGN3378 |
| 2A, 2C-E, 3E-F | MATa, NAM7-GFP::HIS | BY4741 | Yeast GFP CC | yGN3379 |
| 3E-F | MATa, ADR1-GFP::HIS | BY4741 | Yeast GFP CC | yGN3380 |
| 2A, 2C-E, 3E-F | MATa, ACB1-GFP::HIS | BY4741 | Yeast GFP CC | yGN3381 |
| 3E-F | MATa, RPE1-GFP::HIS | BY4741 | Yeast GFP CC | yGN3382 |
| 3E-F | MATa, GRX1-GFP::HIS | BY4741 | Yeast GFP CC | yGN3383 |
| 3E-F | MATa, NCE103-GFP::HIS | BY4741 | Yeast GFP CC | yGN3384 |
| 3E-F | MATa, SGN1-GFP::HIS | BY4741 | Yeast GFP CC | yGN3385 |
| 3E-F | MATa, ASN2-GFP::HIS | BY4741 | Yeast GFP CC | yGN3386 |
| 2A, 2C-E, 3E-F | MATa, ADH6-GFP::HIS | BY4741 | Yeast GFP CC | yGN3387 |
| 2A, 2C-E, 3E-F | MATa, PSP2-GFP::HIS | BY4741 | Yeast GFP CC | yGN3389 |
| 3E-F | MATa, SER2-GFP::HIS | BY4741 | Yeast GFP CC | yGN3390 |
| 4C-F | MATa, LEU2:GFP-Tor1 | W303 | Gift from Robbie Loewith (Backcrossed) [7] | yGN3176 |
| 4E-F | MATa, GFP-Kog1 | W303 | Gift from Robbie Loewith (Backcrossed) [7] | yGN3184 |
| 4E-F | MATa, Lst8-ymNeonGreen:KAN | W303 | This study | yGN3078 |
| 4G | MATa, hog1::KAN | W303 | This study | yGN3406 |
| 2A, 2C-E , 5A-D, 5G-H | MATa, Hog1-GFP::kanMX6 | W303 | This study | yGN3478 |
| 5A-D, 5G-H | MATa, Hog1-GFP-Dbp1LCD::kanMX6 | W303 | This study | yGN3480 |
| 5E-F | MATa, Hog1-GFP::kanMX6, Nup49-mCherry::NAT | W303 | This study | yGN3494 |
| 5E-F | MATa, Hog1-GFP-Dbp1LCD::kanMX6, Nup49-mCherry::NAT | W303 | This study | yGN3495 |

*A list of yeast strains and strain backgrounds used in this study. Yeast GFP CC refers to the yeast GFP clone collection [8] (Thermo).*

**Table S4.**

| Name | Sequence |
|---|---|
| Dbp1-LCD | TTACCTGTCCACACAATCTGCCCTTTCGAAAGATCCCAACGAAAAGAGAGACCACATG<br>GTCCTTCTTGAGTTTGTAACAGCTGCTGGGATTACACATGGCATGGATGAACTATACAA<br>AGGGGCTGGGCTTGGAattCtaAACGCTGACTTGCCTCAGAAGGTAAGCAACTTGAGC<br>ATTAACAATAAAGAAAACGGAGGTGGGGGTGGTAAGTCAAGCTATGTACCGCCCCATC<br>TTCGTTCCCGTGGCAAACCTAGTTTCGAGAGAAGATCTCCCAAACAGAAAGACAAGG<br>TGACAGGGGGAGACTTCTTTCGTCGTGCCGGCAGACAGACAGGGAATAATGGGGGT<br>TTCTTCGGGTTTAGTAAAGAGCGTAATGGTGGGACTTCAGCGAATTACAACCGTAGGG<br>GATCTAGTAACTACAAATCTAGTGGCAACAGATGGGTCAACGGGAAGCACATTCCTGG<br>TCCAAAAAACGCGAAACTGCAAAAAGCTGAGCTATTTGGAGTACATGACGATCCTGAT<br>TATCACAGTTCTGGGATCAAGTTTGATAATTATGATAACATTCCCGTCGATGCAAGTGGT<br>AAGGACGTGCCTGAGCCTATTCTGTAGCCACTTCTAAATAAGCGAATTTCTTATGAT |
| ACT1 FW | CGCTCCTCGTGCTGTCTTCC |
| ACT1 RV | TGGATTGAGCTTCATCACCAACGT |
| STL1 FW | TGGACAGTCCGGTTGGGGTT |
| STL1 RV | TCCGGCGGTTTCAGGGTAGA |
| ALD3 FW | AAGCGCACATGTTTGCTCGC |
| ALD3 RV | TTATCAACGCCGGTGTCGCC |

*A list of oligos used in this study.*
